# Are you my mother?: When host genetics and gut microbiota tell different phylogenetic stories in the Africanized honey bee hybrid (*Apis mellifera scutellata* x sspp.)

**DOI:** 10.1101/2024.10.15.618440

**Authors:** Kilmer Oliveira Soares, Celso José Bruno de Oliveira, Luis Eduardo Martínez Villegas, Priscylla Carvalho Vasconcelos, Adriana Evangelista Rodrigues, Christopher Madden, Vanessa L. Hale

## Abstract

Africanized honey bees (*Apis mellifera scutellata* x sspp.) originated in Brazil through the crossbreeding of African (*A. mellifera scutellata*) and European (*A. mellifera* sspp.) honey bee subspecies. African genes came to dominate in these hybrid honey bees over time. Gut microbiota co-evolve with their hosts and generally reflect host phylogeny. To examine if this was true in Africanized honey bee hybrids, we compared the gut microbiota (16S rRNA) of 3 honey bee subspecies: African, European, and Africanized honey bees. Publicly available sequencing data from five honey bee studies were downloaded from NCBI. European bee samples (n=42) came from United Kingdom, Switzerland, and United States. African bee samples (n=82) came from Kenya. Africanized bee samples (n=10) came from Brazil. Unexpectedly, Africanized honey bee gut microbiota was far more similar to European bees than to African bees despite the closer host genetic relationship between African and Africanized bees. All three subspecies shared similar relative abundances of core taxa. We posit that the similarity in gut microbiota between Africanized and European honey bees arose from the nature of the crossbreeding, and the social / environmental transmission of gut microbiota within hives. Namely, African queens took over European hives. However, the hybrid offspring acquired their gut microbiota from European nurse bees and European hive materials, resulting in the stable transmission of European gut microbiota across generations. Our results provide an intriguing insight into the potential ecological, social, and environmental factors that shape the gut microbiota of the Africanized honey bee hybrid.

**Importance:** Africanized honey bee hybrids originated in Brazil through the crossbreeding of African and European honey bee subspecies. In this study, we examined the gut microbiota of all 3 honey bee subspecies (African, European, Africanized). A few core microbiota were shared across all subspecies. Interestingly, while African honey bee genes dominated in the Africanized honey bee hybrids, their gut microbial composition was most similar to European bees. This is likely related to the way these bees were crossbred – with African queens taking over European hives, while gut microbial inoculation of hybrids came from European nurse bees and European hive matierals. Gut microbiota are critical to honey bee health, and studying the gut microbiota of closely related honey bee subspecies helps understand the factors that influence gut microbial composition. This is important for our broader understanding of honey bee health, conservation, and evolution.

## INTRODUCTION

Honey bees have spread throughout the world via migration, hybridization, and human importation (CALFEE *et al*., 2020; LEGESSE; GETU, 2022; MORTENSEN; SCHMEHL; ELLIS, 2013). Thirty-three genetically distinct *Apis mellifera* honey bee subspecies have been identified and are geographically distributed across three main regions: Africa (11 subspecies), Western Asia and the Middle East (9 subspecies), and Europe (13 subspecies) (ILYASOV *et al*., 2020). Brazil is home to a hybid honey bee subspecies (*Apis mellifera scutellata* x sspp.) derived from a cross between African and European subspecies. European honey bees (*Apis mellifera* sspp.) were first introduced to Brazil in 1839 by priest Antonio Pinto Carneiro who imported bees from Portugal and Spain (DE QUEIROZ ROLIM *et al*., 2018). Later, between 1845 and 1880, German and Italian immigrants introduced other Eurpoean subspecies (*A. mellifera* sspp.) to Brazil (DE QUEIROZ ROLIM *et al*., 2018). In 1956, Professor Warwick Estevam Kerr brought 26 African honey bee queens (*Apis mellifera scutellata)* from Tanzania and South Africa to Brazil. These queens escaped from their apiary and attacked and replaced the queens in the European bee colonies already established in Brazil. The African queens then mated with European drones ultimately establishing a new hybrid subspecies (SCOTT SCHNEIDER; DEGRANDI-HOFFMAN; SMITH, 2004). This was the origin of the Africanized honey bee (*Apis mellifera scutellata* x sspp.) (MICHAEL; J. D.; NALEN, 2015) (**Figure 1**).

**Figure 1.**
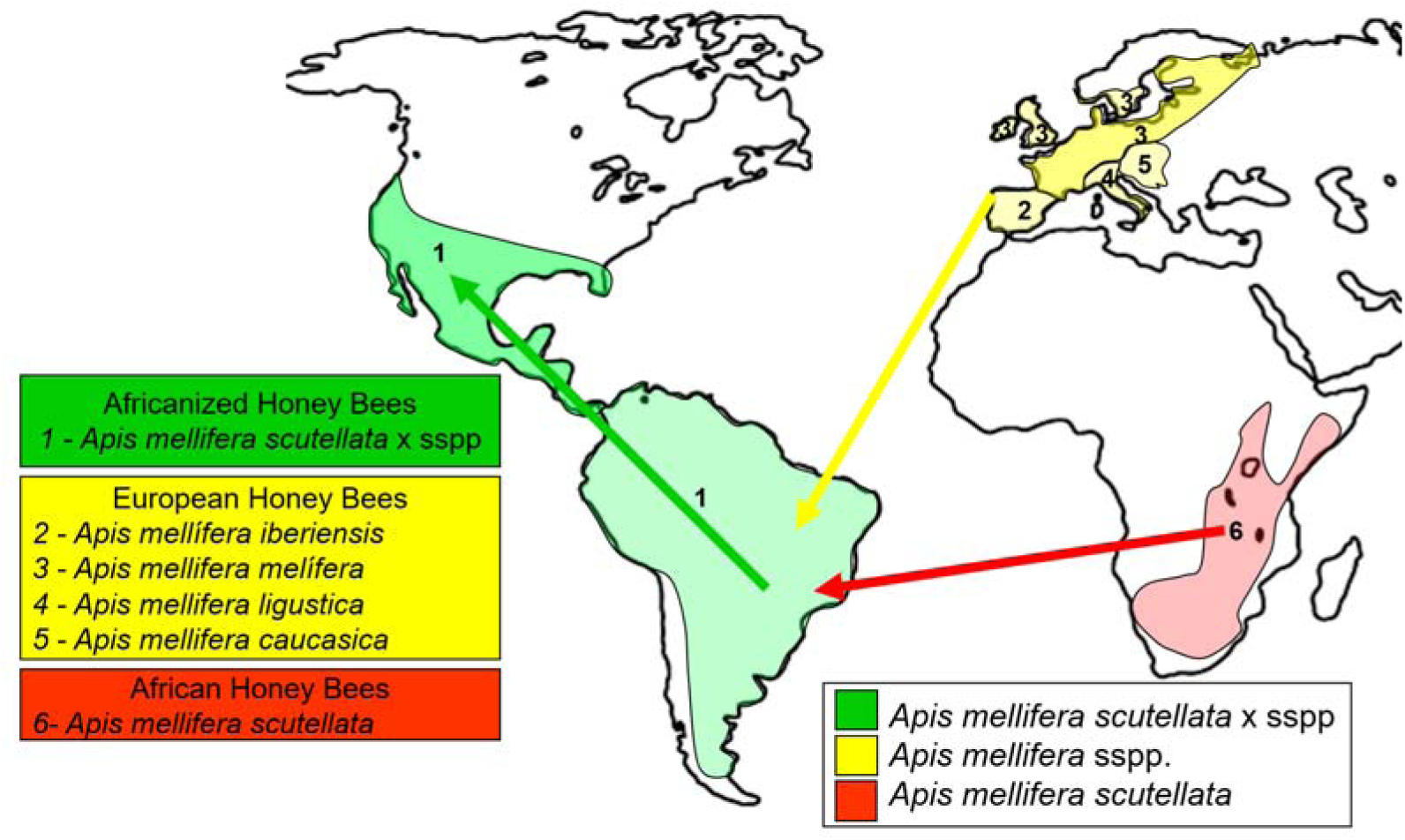
Honey bee introductions, hybridization and expansion. Yellow route: European honey bees (*Apis mellifera* sspp.) were introduced in Brazil between 1839 and 1880 (De Queiroz Rolim *et al*., 2018). Red route: African honey bee queens (*Apis mellifera scutellata*) were introduced in Brazil in 1956 (Sandford, 2004); Green route: Natural expansion of Africanized honey bees (*Apis mellifera scutellata* x sspp) across Americas (adapted from (Pucca et al., 2019).

Honey bee subspecies exhibit differing biological characteristics (e.g. pathogen resistance, heat tolerance, behavior, resource use) that have supported their adaptation and expansion across varying environments, climates, and ecological pressures around the globe. African honey bee subspecies, for example, exhibit rapid colony growth, increased resistance to Varroa mites and pathogenic viruses, and improved survival in neotropical environments as compared to European honey bee subspecies (TIBATÁ *et al*., 2021). These same traits proved beneficial in Brazil, resulting in the dominance of African-bee derived genes (50– 90%) in the Africanized honey bee hybrid, and facilitating the spread of this hybrid subspecies (*A. mellifera scutellata* x sspp) across the the Americas (**Figures 1, 2**). (CALFEE *et al*., 2020; CHAPMAN *et al*., 2015; COLLET *et al*., 2006). Although many phenotypic and genotypic differences have been characterized between African, European, and Africanized honey bees, one biological characteristic that has not been well explored in relation to honey bee phylogeny and hybridization is the gut microbiota.

**Figure 2:**
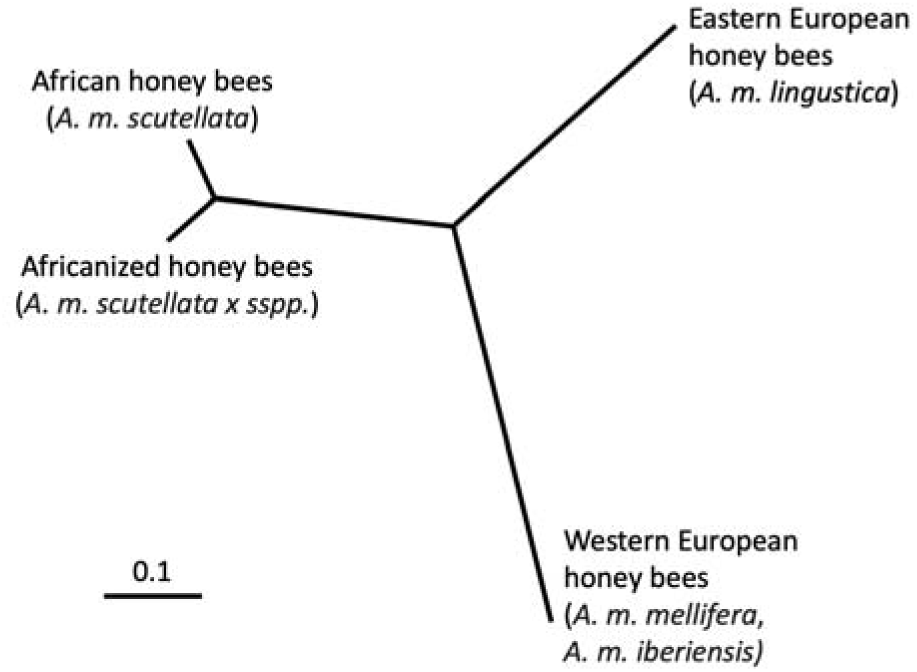
Genetic distance between African, European, and Africanized honey bee populations. Distances based on 95 single nucelotide polymorphisms (SNPs) displayed in an unrooted UPGMA tree. Adapted from Chapman et al. 2015.

Importantly, gut microbiota are critical to host health and the adaptation of hosts to new environments (KEŠNEROVÁ *et al*., 2020; ZHANG *et al*., 2024). In fact, previous studies have already determined that honey bee gut microbes play a key role in nutrient degradation (LEE *et al*., 2015), bacterial and fungal pathogen defense (ENGEL *et al*., 2016; MILLER; SMITH; NEWTON, 2021; MOTTA; RAYMANN; MORAN, 2018), pesticide tolerance (WU *et al*., 2020), and behavior (TESEO *et al*., 2019; VERNIER *et al*., 2020). Numerous prior studies have also demonstrated links between host phylogeny and the gut microbiome (BRANSTETTER *et al*., 2017; ENGEL; MORAN, 2013; KWONG *et al*., 2017; SARTON-LOHÉAC *et al*., 2023). Understanding the relationship between host phylogeny, hybridization, and the gut microbiota of these three honey bee subspecies (African, European, Africanized) is of significant importance as it can provides insights into mechanisms of adaptation and evolution in honey bees. In this study, we compared gut microbiota of African, European, and Africanized honey bee subspecies. We hypothesized that Africanized honey bee gut microbiota (*A. mellifera scutellata* x sspp) would represent a hybrid between African and European bee gut microbiota, and that Africanized bee gut microbiota would align with host phylogeny and be more closely related to African bee gut microbiota.

## 1. METHODS

### 1.1. STUDY DESIGN

This study included publicly available 16S rRNA data from five previously published studies on honey bees (**Figure 3, Supplemental Table 1**). Data was downloaded from NCBI’s short read archive (SRA). Sequencing platforms, primers, and extraction methods used in each study are listed in Supplemental Table 1. All studies sampled free-ranging adult workers bees under natural field conditions. We exclusively used “control bee” samples if the study involved treatment and control groups. European honey bee samples came from: Austin, Texas – United States (n=15) (MOTTA; RAYMANN; MORAN, 2018); Sussex, United Kingdom (n=15) (JONES *et al*., 2018); and Laussane, Switzerland (n=14) (KEŠNEROVÁ *et al*., 2020). African honey bee samples came from four sites in Kenya: Kakamega (n=17), Kilifi-Coast (n=22), Kwale-Coast (n=20), and Nairobi (n=23) (TOLA *et al*., 2020). Africanized honey bee samples came from two municipalities of the State of Paraíba – Brazil: Areia (n=5) (SOARES *et al*., 2021) and São João do Cariri (n=5). The honey bee samples from São João do Cariri (7°22’55.7“S 36°31’40.9”W; Altitude 458m) were obtained for the purpose of this comparative study, according to the methods described below.

**Figure 3.**
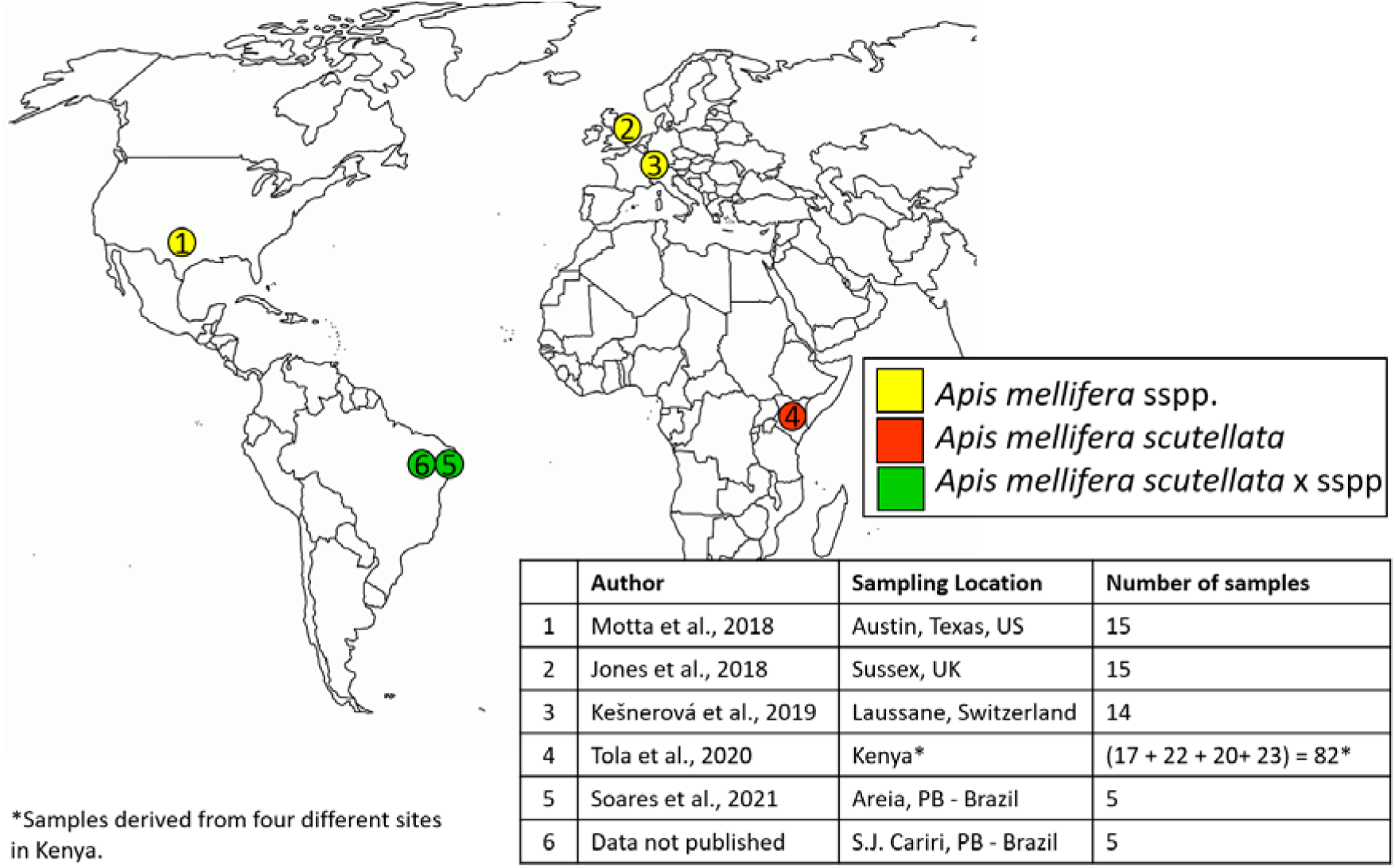
Sampling locations for European, African, and Africanized honey bees.

### 1.2. SAMPLING, DNA EXTRACTION, LIBRARY PREPARATION, AND SEQUENCING

For Africanized honey bees from São João do Cariri-PB, 100 adult bees were collected from hive frames, placed in sterile tubes containing 70% ethanol, and transported to the lab (Laboratory for the Evaluation of Products of Animal Origin - LAPOA, Areia PB) and stored at –20°C until extraction. All procedures performed were approved by the Biodiversity Authorization and Information System—SISBIO (Protocol #: 71750-1, approved on 09/19/2019). Honey bee dissection, DNA extraction, and library preparation were conducted as described previously (SOARES *et al*., 2021). Each sample (n=5) represented the pooled abdominal contents of 20 bees. Paired-end sequencing of the V3-V4 region (Primers: 341F, 805R) was performed on an Illumina MiSeq with a V2 kit (2 × 250 cycles).

### 1.3. SEQUENCE DATA PROCESSING AND STATISTICAL ANALYSIS

Raw paired-end sequences were downloaded for each study separately and then merged using the command “qiime feature-table merge”. Sequences were then demultiplexed, filtered, denoised, and truncated to a length of 248 base pairs using QIIME 2-2020.2 (Bolyen *et al*., 2019) (See **Appendix 1** for QIIME Scripts). According to Liu *et al*. (2020), V3-V4 amplicon analysis is compatible with V4 amplicon analysis after trimming to the same region (**Appendix 1 – Script 1**). Chimeric sequences were removed, and DADA2 was used to parse sequences into Amplicon Sequence Variants (ASVs) (CALLAHAN *et al*., 2016). Sequences were then aligned using “qiime fragment-insertion sepp” for phylogenetic analysis (MATSEN *et al*., 2012). Taxonomic composition of the samples was determined with a pretrained naive Bayes classifier with a 99% sequence similarity threshold for V3-V4 reference sequences (SILVA-132-99-nb-classifier.qza) and the “qiime feature-classifier classify-sklearn”. Reads identified as chloroplasts, mitochondria, unassigned and eukaryota were removed from all samples.

Microbial composition and diversity were analyzed in QIIME 2-2020.2 (Bolyen *et al*., 2019). Microbial composition (beta diversity) was compared between honey bee subspecies (European, African, Africanized) using PERMANOVAs based on Bray-Curtis, Jaccard, and Weighted and Unweighted Unifrac distances (ANDERSON, Marti J, 2001) Microbial communities were visualized using Principle Coordinate Analysis (PCoA) and the Emperor plugin 2020.2.0 (VÁZQUEZ-BAEZA *et al*., 2017). Microbial diversity (alpha diversity) was assessed using observed ASVs, the Shannon diversity index (richness and abundance), Faith’s PD (phylogenetic diversity), and Pielou’s (evenness) diversity index (Sampson *et al*., 2016). We assessed for normality using Shapiro-Wilk tests; then, as appropriate used ANOVAs or Tukey or Kruskal-Wallis Tests at 5% probability to compare microbial diversity between groups using R version 4.1.0 (RIPLEY, 2001).

Differentially abundant taxa were identified using an analysis of composition of microbiomes (ANCOM) (Mandal *et al*., 2015). We also identified core taxa, defined here as taxa present in 90% of the samples using the command “qiime feature-table core-features”. Both analyses were performed at the genus level. The relative abundances of core microbes were compared by honey bee subspecies using a one-way ANOVA followed by a pairwise Kruskal-Wallis Rank Sum Test **(Appendix 1-Script 2)**. To assess the strength of the relationship between the occurance of specific taxa and honey bee subspecies, a multi-level pattern analysis was performed in R (multipatt function, package Indicspecies v.1.7.12, function “r.g”, α=0.5). This analysis, assigns a test statistic, or ‘indicator value’ to each ASV in each group (DE CÁCERES *et al*., 2011; DE CÁCERES; LEGENDRE, 2009) **(Appendix 1 -Script 3)**.

## 2. RESULTS

This study included samples from three honey bee subspecies derived from six locations worldwide: African honey bees (*A. mellifera scutellata*, n=82), European honey bees (*A. mellifera* sspp., n = 44), and Africanized honey bees (*A. mellifera scutellata x sspp*, n=10) (**Figure 3**, **Supplemental Table 1**).

### 3.1 MICROBIAL COMPOSITION AND DIVERSITY BY HONEY BEE SUBSPECIES AND PROVENANCE

We obtained a total of 919,168 raw reads across all samples, with an average of 6,565 reads per sample (range: 1,015 to 26,958 reads). Overall honey bee gut microbial composition differed significantly by subspecies (PERMANOVA: Weighted Unifrac R^2^ = 0.4178, *p*-value = 0.001; Unweighted Unifrac R^2^ = 0.3062, *p*-value = 0.001, **Figure 4, Supplemental Figure 1**). However, based on pairwise comparisons (Weighted UniFrac), European and Africanized honey bees did not differ significantly while African bees differed significantly from both European and Africanized bees (**Figure 4d**, Weighted Unifrac Pairwise PERMANOVAs: Africanized vs. European: *p* = 0.136; Africanized vs. African: *p* = 0.001, African vs. European: *p* = 0.001; Unweighted Unifrac Pairwise PERMANOVAs: Africanized vs. European p = 0.001; European vs. African: p = 0.001; African vs. Africanized p = 0.001). Microbial diversity also differed significantly between subspecies (Kruskal-Wallis: Shannon Index *p* = 2.2e-16; Faith’s PD *p* = 2.2e-16; Evenness Pielou *p* = 1.421e-08; Observed Features *p* = 2.2e-16, **Figure 5**). Africanized honey bees exhibited a microbial diversity (Shannon diversity index) significantly greater than European bees but lesser than African bees. In terms of phylogenetic diversity (Faith’s PD) and evenness (Pileou’s index), Africanized honey bees were comparable to European bees, while in terms of microbial richness, Africanized honey bees were comparable to African bees.

**Figure 4.**
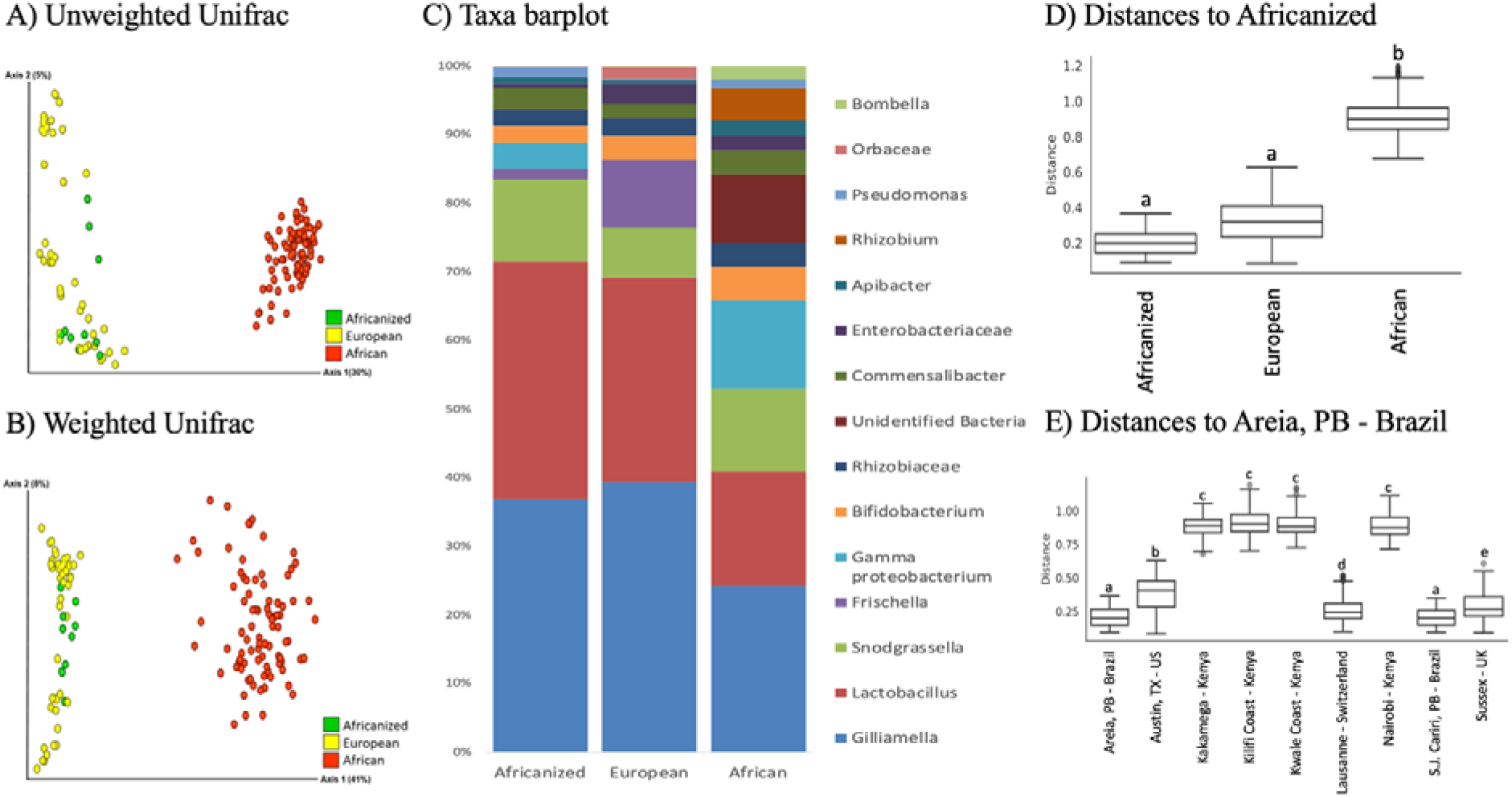
Gut microbial composition of European, African, and Africanized honey bees. Overall honey bee gut microbial composition differed significantly between all three honey bee subspecies based on A) Unweighted UniFrac and B) Weighted UniFrac metrics. C) Taxa barplot of all microbial taxa found at >1% relative abundance and averaged across all samples from each honey bee subspecies. D) Pairwise comparisons (Weighted UniFrac) between honey bee subspecies (Weighted UniFrac pairwise PERMANOVAs: Africanized vs. European *p* = 0.136; European vs. African *p* = 0.001; African vs. Africanized *p* = 0.001). E) Pairwise comparisons (Weighted UniFrac) by honey bee provenance (Weighted UniFrac PERMANOVA *p* = 0.001; see **Supplemental Table 3** for pairwise p values.)

**Figure 5.**
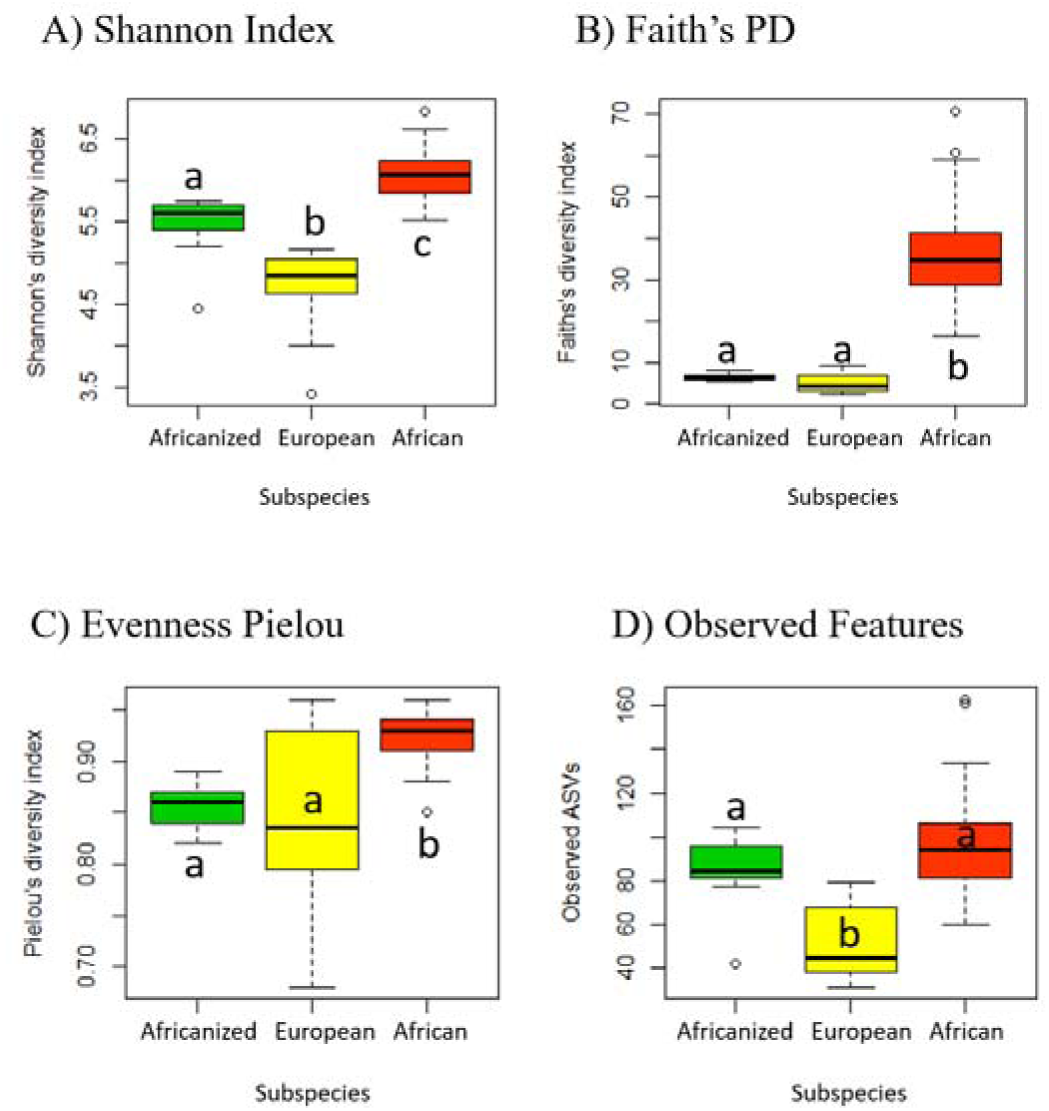
Gut microbial diversity of European, African, and Africanized honey bees. A) Shannon Index, B) Faith’s PD, C) Pielou’s Evenness, D) Observed Features. Box plots show outliers, first and third quartiles (lower and upper edges), and highest, lowest, and median values (horizontal black dash). (Kruskal-Wallis: Shannon Index *p* = 2.2e-16; Faith’s PD *p* = 2.2e-16; Evenness Pielou *p* = 1.421e-08; Observed Features *p* = 2.2e-16). Significant differences (*p* ≤ 0.05 Tukey’s test)) are denoted with different letters over each box plot.

In an effort to parse the effects of provenance (biogeography) versus host (subspecies), we then examined gut microbial composition by honey bee provenance. Gut microbial communities from all nine sampling locations around the world differed significantly, except the four locations in Kenya compared to each other and the two locations in Brazil compared to each other (**Figure 4e**, **Supplemental Figure 2, Supplemental Tables 2, 3**; Weighted UniFrac PERMANOVA *p* = 0.001). Notably, the two locations in Brazil differed signicantly on weighted but not unweighted UniFrac metrics indicating that gut microbial taxa were similar across locations, but there were differences in relative abundances of these taxa by sampling location.

### 3.2 CORE MICROBIOTA AND DIFFERENTIALLY ABUNDANT MICROBAL TAXA

A core microbiota analysis identified three taxa (genera) that were present in 90% of the samples across all honey bee subspecies. These included: *Lactobacillus, Snodgrassella*, and *Gilliamella* (**Figure 6**). These taxa accounted for 15% of all genera in the dataset. There were no significant differences in abundances of these taxa across subspecies (Kruskal-Wallis all p > 0.4).

**Figure 6.**
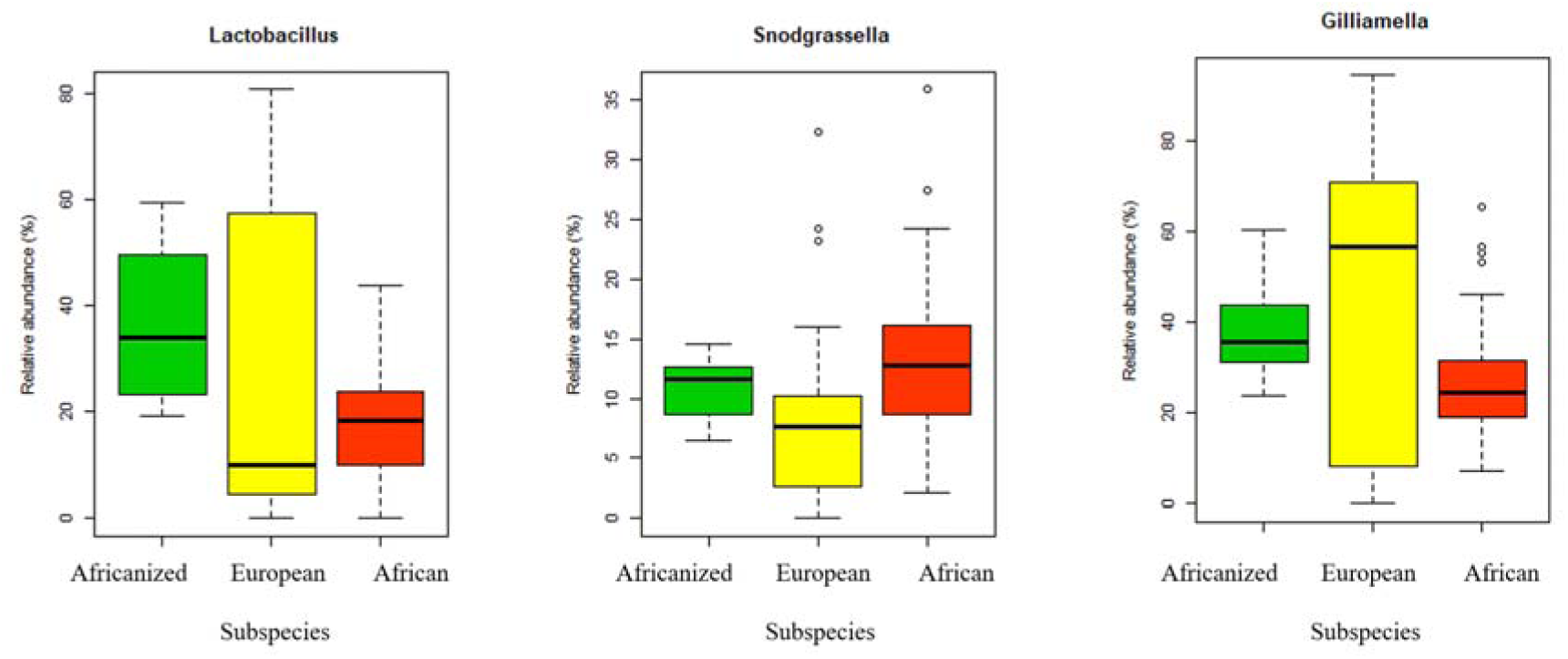
Relative abundances of core microbiota (genera) that were present in 90% of the samples across all subspecies. A) Lactobacillus, B) Snodgrassella, and C) Gilliamella. Box plots show outliers, first and third quartiles, and highest, lowest, and median values. None of these taxa differed significantly in abundance by honey bee subspecies (Kruskal-Wallis all p > 0.4).

We further identified nine differentially abundant taxa (genera) by honey bee subspecies including: taxa in the families Orbaceae, Rhizobiaceae (Allorhizobium-Neorhizobium-Pararhizobium-Rhizobium) and Enterobacteriaceae, taxa from the genera *Frischella, Pantoea, Bombella, Gilliamella,* a taxa identified as a Gammaproteobacterium, and an unidentified Bacteria (**Figure 7, Supplemental Table 4**). Africanized and African bees shared similar relative abundances of the Gammaproteobacterium, and the Enterobacteriaceae, *Pantoea*, *Bombella*, and *Orbaceae* taxa. Africanized and European bees shared similar relative abundances of: *Gilliamella*, *Fischella*, *Pantoea*, *Bombella*, the unidentified Bacteria, and the Enterobacteriacea and Rhizobiaceae taxa. African and European bees differed significantly in relative abundances of all these taxa.

**Figure 7.**
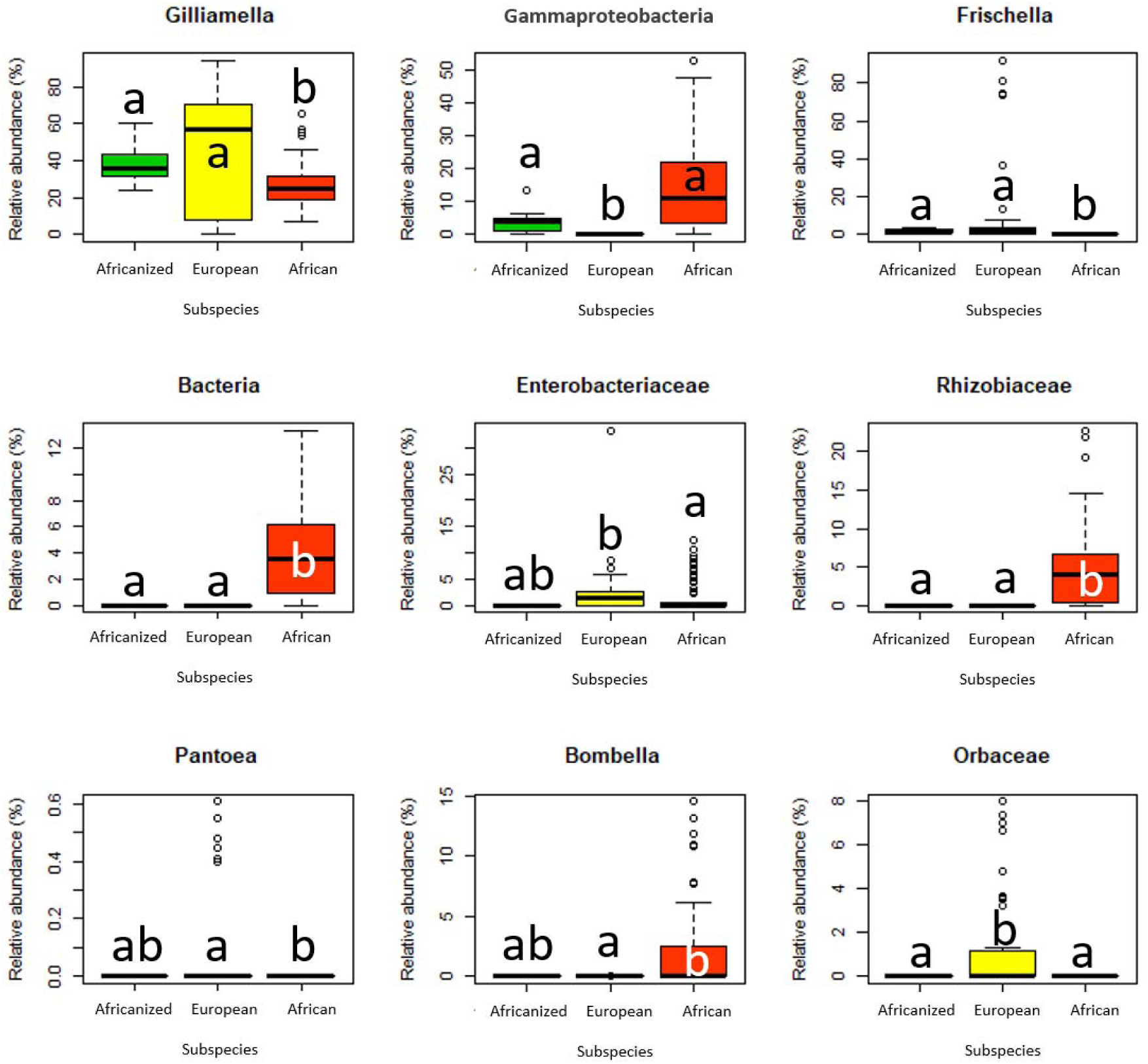
Relative abundances of differentially abundant genera (ANCOM) by honey bee subspecies. Box plot shows outliers, first and third quartiles, and highest, lowest, and median values. Stastitical differences (Tukey’s test p ≤ 0.05) are denoted by different letters over each box plot.

### 3.3 MICROBIAL INDICATOR SPECIES

To determine which microbial taxa distinguished each honey bee subspecies or subspecies pair, we performed an indicator species analysis (R package Indicspecies v.1.7.12). Taxa with an indicator value of >0.4 and *p* ≤ 0.05 were considered microbial indicator species. Species within the Alphaproteobacteria class (e.g., *Bombella*) were key indicators of African bees while Gammaproteobacteria species (e.g., *Pantoea*, *Frischella*) were indicators in European bees. An uncultured Gammaproteobacterium was shared between African and Africanized honey bees while a *Gilliamella* species. was shared between European and Africanized honey bees (**Supplemental Table 5).**

## 3. DISCUSSION

In this study, we compared gut microbiota of three honey bee subspecies (African, European, and Africanized) and found significant differences in microbial composition and diversity between subspecies and, to a lesser degree, by honey bee provenance. We hypothesized that Africanized honey bees, which are a hybrid between African and European honey bees, would have gut microbial communities that shared features with both African and European bees. Our results supported this hypothesis. We further hypothesized that Africanized honey bee gut microbiota would be most similar to African honey bee gut microbiota as these two subspecies are most closely related phylogenetically. However, our results did not support this hypothesis: Instead, we found that Africanized and European honey bees had much more similar gut microbial communities than African and Africanized bees despite being more phylogenetically distant (less genetically related) based on host (honey bee) genes. We explore potential explanations for this finding below.

### 4.1 AFRICANIZED AND EUROPEAN HONEY BEES EXHIBITED HIGHLY SIMILAR GUT MICROBIOTA DESPITE PHYLOGENETIC DISTANCE BETWEEN HOSTS

Africanized and European honey bees exhibited similar overall gut microbial composition and dominant taxa (Weighted UniFrac), similar phylogenetic diversity (Faith’s PD) and evenness (Pileou’s), and similar abundances of many common honey bee taxa (e.g., *Gilliamella, Frichella, Bombella*, and taxa in the Rhizobiaceae family), with a *Gilliamella* species notably serving as an “indicator species” found at ∼35-60% relative abundance in European and Africanized bees and only up to ∼25% relative abundance in African bees. We posit that the similarity between Africanized and European bee gut microbiota may be due to honey bee hive dynamics and sociality in shaping microbiota transmission and acquisition.

Specifically, honey bees acquire gut microbiota through contact with nurse bees, fecal material, and hive components (comb, honey, bee bread) (ENGEL *et al*., 2016; POWELL *et al*., 2014). Gram-positive bacteria, such as *Lactobacillus* or *Bifidobacteria*, are typically acquired through contact with hive surfaces and nurse bees, including via trophyllaxis (nutrient exchange by mouth from nurse to newly eclosed adult bees) (ANDERSON, Kirk E. *et al*., 2022; POWELL *et al*., 2014; VÁSQUEZ *et al*., 2012). Acquisition of gram-negative bacteria such as *Gilliamella, Frischella,* and *Snodgrassella* relies on contact with fresh fecal material in the hive, or contact with nurse bees (not including trophyllaxis) (POWELL *et al*., 2014). This social and environmental gut microbial transmission/acquisition process ultimately leads to stable coadapted gut microbial communities across generations in adult honey bees (ELLEGAARD; ENGEL, 2019; KOWALLIK; MIKHEYEV, 2021; KWONG *et al*., 2017; KWONG; MORAN, 2016; MORAN *et al*., 2012; POWELL *et al*., 2014).

When African queens were accidentally released in Brazil and took over existing European honey bee hives by killing the European queens, although the African queens maintained their own distinct gut microbiota, the nurse bees, and hive materials were all from the original European bee hives. Thus, even though the newly emerging Africanized bees were genetically more closely related to the African queen, these hybrids were still acquiring gut microbiota via contact with European nurse bees, fecal material, and hive components.

This idea is supported by the Tarpy et al. (2015) study demonstrating that the queen gut microbiome is distinct from and not transmitted to offspring within a hive, and that there are only modest changes to worker bee microbiota (in terms of relative abundances) before and after queen replacement. Ultimately, the European bee gut microbiota may have also provided fitness benefits to the Africanized hybrids, as European bees had been present in and adapting to the Brazilian climate and environment for over 100 years before African bees were introduced.

### 4.2 AFRICANIZED AND AFRICAN HONEY BEES MAINTAINED FEW SIMILARITIES IN GUT MICROBIOTA DESPITE PHYLOGENETIC CLOSENESS BETWEEN HOSTS

African and Africanized honey bees shared fewer similarities in gut microbiota as compared to European and Africanized bees. African honey bee gut microbiota was highly distinct from Africanized bees and generally more diverse (Shannon Index, Faith’s PD, Pileou’s Evennnes). This could be due to minimal transmission of gut microbiota from African queens to Africanized hybrids, or to differing diets, dietary diversity, environments, or environmental pressures faced by African bees as compared to European and Africanized bees (BROWN; MAYES; BHATTA, 2016; KWONG *et al*., 2017; KWONG; MANCENIDO; MORAN, 2017). A few taxa, including a Gammaproteobacteria and a taxa in the Orbaceae family, were found at similar abundances in African and Africanized bees. Gammaproteobacteria abundances have been reported to increase in response to stressors including parasite and pesticide exposures (PARIS et al., 2020). It is feasible that honey bees in Africa and Brazil faced some similar stressors and that increased Gammaproteobacteria abundances represent a response to these stressors. It is also possible that Gammaproteobacteria provide fitness benefits to honey bees in both Africa and Brazil (e.g. pathogen resistance, rapid colony growth, adaptability to new environments) (POWELL *et al*., 2016; TIBATÁ *et al*., 2021). Alternately, these taxa may be particularly well suited to social transmission (e.g., from queen to workers to offspring) (TARPY; MATTILA; NEWTON, 2015), or have a microbiome-host genetic associations linked to African honey bee genes.

### 4.3 CORE MICROBIOTA ACROSS HONEY BEE SUBSPECIES

*Lactobacillus*, *Snodgrassella*, and *Gilliamella* were identified as core microbiota present in 90% of the samples and found across all honey bee subspecies. Other studies report similar findings in terms of core microbiota (KWONG *et al*., 2017). Notably, *Bifidobacterium* is also considered a core microbe in honey bees (KWONG *et al*., 2017), but was only present in 80% of the samples in our study and therefore did not meet our “core” threshold. Fourteen (out of 82) samples from African bees lacked detectable *Bifidobacterium,* which may been an artifact of sample preservation, extraction, or sequencing as *Bifidobacterium* is generally found at lower abundances in the honey bee gut (MORAN *et al*., 2012).

### 4.4 GUT MICROBIOTA AND HONEY BEE PROVENANCE

We observed the most pronounced differences in honey bee gut microbiota by host phylogeny (subspecies), but we also observed differences in gut microbiota based on honey bee provenance. Provenenance is linked with environment and diet, and multiple studies have reported differences in gut microbiota based on provenance in various animal species including honey bees (KEŠNEROVÁ *et al*., 2020; MINICH *et al*., 2021; MORAN *et al*., 2012; SOARES *et al*., [*s. d.*]). Notably, as our study employed publicly available data from other studies, there were differences in sample preservation, extraction, and sequencing methods that potentially confounded these analyses (**Supplemental Table 1**). This was particularly true across European bee samples / studies. However, all African honey bee samples (from four sampling locations across Kenya, central to coast) were collected, extracted, and sequenced in the same manner, and when these technical variables were controlled for, no differences were detected in gut microbiota by sampling location in African honey bees. These results support the original findings of the Tola et al. 2020 study. Interestingly, the honey bee samples from the two locations in Brazil were also preserved, extracted and sequenced in the same way, but *did* differ significantly in microbial taxonomic abundances (Weighted UniFrac), however not in unweighted taxonomic profiles (**Supplemental Tables 2, 3**). Taken together, this suggests that provenance does impact honey bee gut microbiota but that host phylogeny may play a larger role in shaping bee gut microbial communities.

## 5. CONCLUSION

Our study provides key insights into potential factors that have shaped Africanized honey bee gut microbiota co-evolution. Namely, our results suggest that the introduction of African queens to European hives, and the social / environmental transmission of gut microbiota within these hives lead to an Africanized honey bee hybrid that is phylogenetically more African while hosting a gut microbiota that is more European. Provenance can also play a role in shaping honey bee gut microbiota, although to a lesser degree. Deeper examination of microbial phylogenies between honey bee subspecies, and evaluation of honey bee fitness across environments with differing microbial strains could further elucidate the mechanisms underlying these patterns. Examining the complex relationships between honey bees and their gut microbiota is critical for understanding honey bee evolution, for conservation efforts and the preservation of host and microbial genetic diversity in honey bee populations, for beekeeping practices, and ultimately for protecting the health and sustainability of these vital pollinators.

## Data Availability Statement

The datasets presented in this study can be found in online repositories. The names of the repository/repositories and accession number(s) can be found below: NCBI (accession: PRJNA732391).

## Supporting information

Supplemental material: Scripts from R and Qiime2

## SUPPLEMENTAL MATERIAL

### Supplemental figures

**Supplemental Figure 1.**
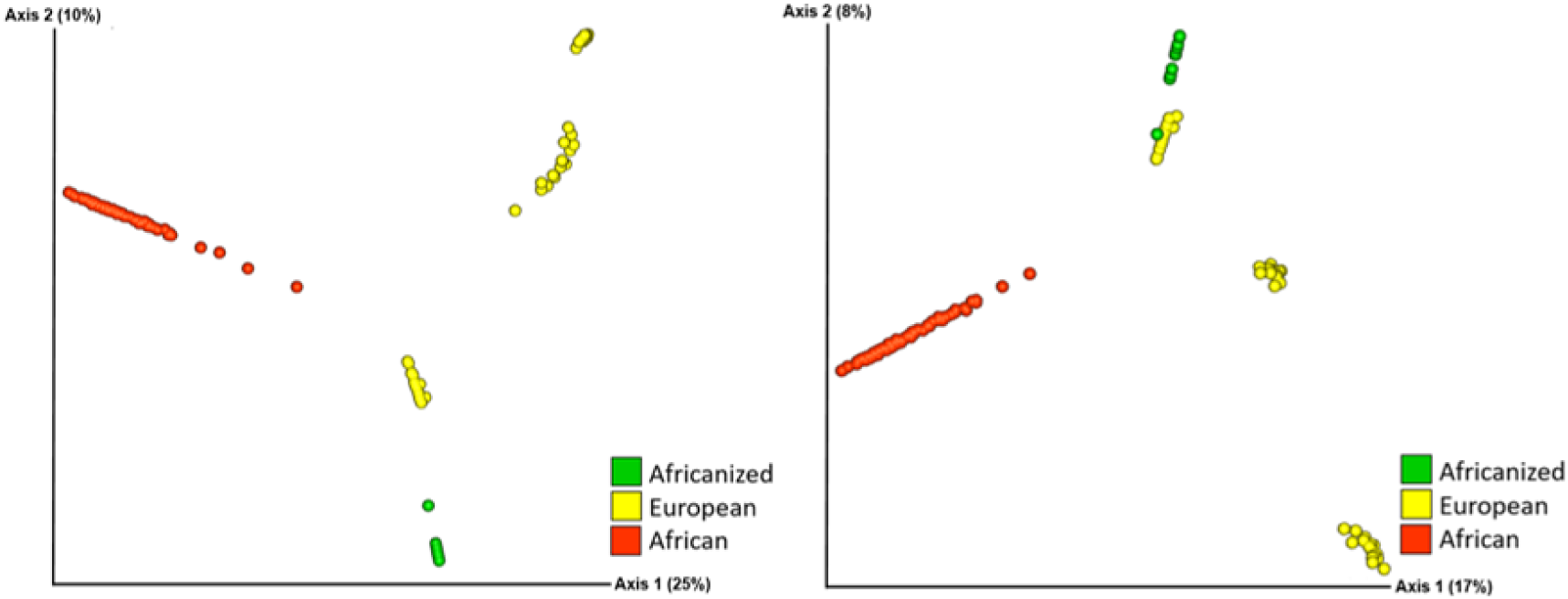
Gut microbial composition of European, African, and Africanized honey bees. Overall honey bee gut microbial composition differed significantly between all three honey bee subspecies based on A) Bray-Curtis and B) Jaccard metrics. (PERMANOVA: Bray Curtis R^2^ = 0.3186, *p* = 0.001; Jaccard R^2^ = 0.2119, *p* = 0.001).

**Supplemental Figure 2.**
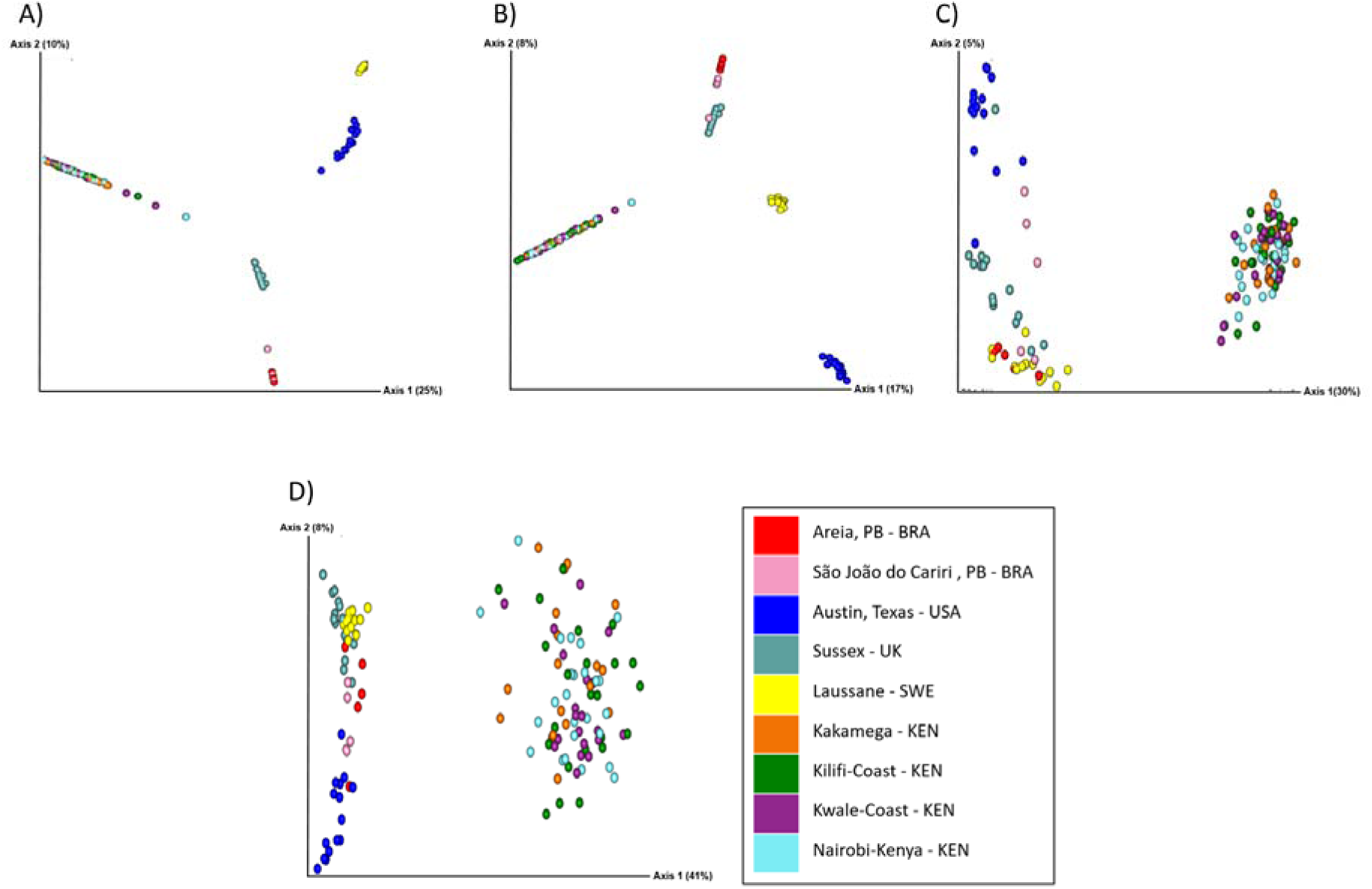
Gut microbial composition of bees based on provenance. Honey bee gut microbial composition differed significantly between all sampling locations on A) Bray-Curtis, B) Jaccard, C) Unweighted Unifrac and D) Weighted Unifrac metrics (PERMANOVA: Bray Curtis, *p* = 0.001; Jaccard, *p* = 0.001, Unweighted Unifrac *p* = 0.001, Weighted Unifrac, *p* = 0.001).

### Supplemental Tables

**Supplemental Table 1.**
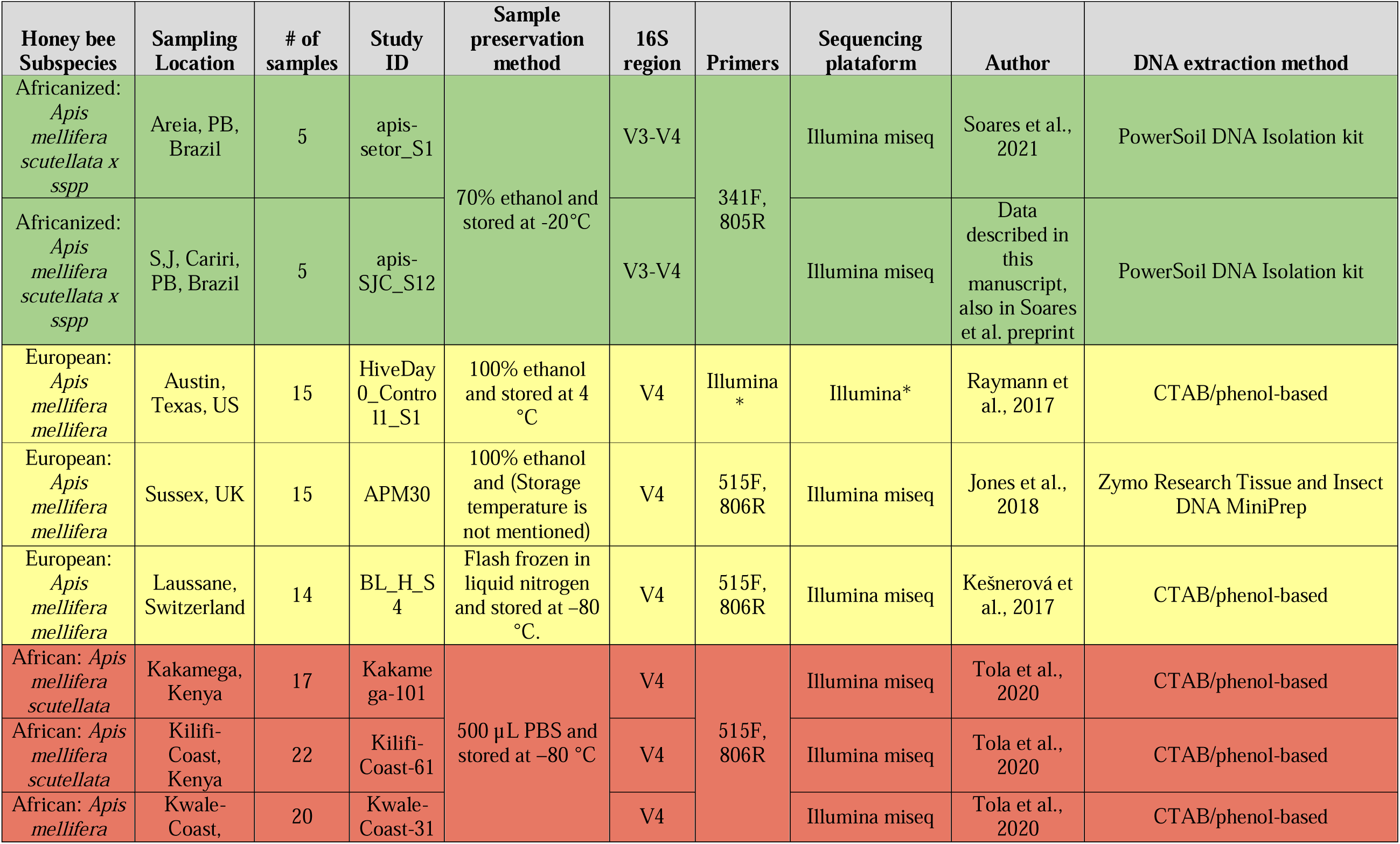
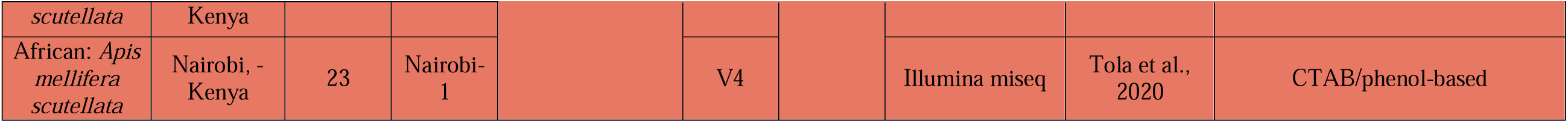
Sampling locations and study information for all honey bee samples included in this study.

**Supplemental Table 2.** Pairwise comparisons of gut microbial communities by sampling location. There was a significant diference in gut microbial composition (Unweighted UniFrac) between all sampling locations except the four locations within Kenya compared to each other.

| Group 1 | Group 2 | Sample size | Permutations | pseudo-F | p-value | q-value |
| --- | --- | --- | --- | --- | --- | --- |
| Areia, PB - Brazil | Austin, Texas - US | 20 | 999 | 18.56 | 0.001 | 0.001 |
| Areia, PB - Brazil | Kakamega-Kenya | 23 | 999 | 8.49 | 0.001 | 0.001 |
| Areia, PB - Brazil | Kilifi-Coast-Kenya | 28 | 999 | 8.39 | 0.001 | 0.001 |
| Areia, PB - Brazil | Kwale-Coast-Kenya | 25 | 999 | 8.12 | 0.001 | 0.001 |
| Areia, PB - Brazil | Laussane, Switzerland | 19 | 999 | 3.55 | 0.002 | 0.002 |
| Areia, PB - Brazil | Nairobi-Kenya | 30 | 999 | 8.36 | 0.001 | 0.001 |
| Areia, PB - Brazil | S.J. Cariri , PB - Brazil | 10 | 999 | 2.24 | 0.011 | 0.013 |
| Areia, PB - Brazil | Sussex, UK | 20 | 999 | 4.94 | 0.001 | 0.001 |
| Austin, Texas - US | Kakamega-Kenya | 33 | 999 | 26.54 | 0.001 | 0.001 |
| Austin, Texas - US | Kilifi-Coast-Kenya | 38 | 999 | 26.51 | 0.001 | 0.001 |
| Austin, Texas - US | Kwale-Coast-Kenya | 35 | 999 | 25.94 | 0.001 | 0.001 |
| Austin, Texas - US | Laussane, Switzerland | 29 | 999 | 17.98 | 0.001 | 0.001 |
| Austin, Texas - US | Nairobi-Kenya | 40 | 999 | 26.91 | 0.001 | 0.001 |
| Austin, Texas - US | S.J. Cariri , PB - Brazil | 20 | 999 | 8.68 | 0.001 | 0.001 |
| Austin, Texas - US | Sussex, UK | 30 | 999 | 13.60 | 0.001 | 0.001 |
| Kakamega-Kenya | Kilifi-Coast-Kenya | 41 | 999 | 0.81 | 0.889 | 0.909 |
| Kakamega-Kenya | Kwale-Coast-Kenya | 38 | 999 | 0.99 | 0.481 | 0.525 |
| Kakamega-Kenya | Laussane, Switzerland | 32 | 999 | 15.17 | 0.001 | 0.001 |
| Kakamega-Kenya | Nairobi-Kenya | 43 | 999 | 1.10 | 0.231 | 0.268 |
| Kakamega-Kenya | S.J. Cariri , PB - Brazil | 23 | 999 | 6.48 | 0.001 | 0.001 |
| Kakamega-Kenya | Sussex, UK | 33 | 999 | 19.22 | 0.001 | 0.001 |
| Kilifi-Coast-Kenya | Kwale-Coast-Kenya | 43 | 999 | 0.82 | 0.909 | 0.909 |
| Kilifi-Coast-Kenya | Laussane, Switzerland | 37 | 999 | 15.73 | 0.001 | 0.001 |
| Kilifi-Coast-Kenya | Nairobi-Kenya | 48 | 999 | 0.96 | 0.572 | 0.606 |
| Kilifi-Coast-Kenya | S.J. Cariri , PB - Brazil | 28 | 999 | 6.54 | 0.001 | 0.001 |
| Kilifi-Coast-Kenya | Sussex, UK | 38 | 999 | 19.84 | 0.001 | 0.001 |
| Kwale-Coast-Kenya | Laussane, Switzerland | 34 | 999 | 14.81 | 0.001 | 0.001 |
| Kwale-Coast-Kenya | Nairobi-Kenya | 45 | 999 | 1.11 | 0.246 | 0.277 |
| Kwale-Coast-Kenya | S.J. Cariri , PB - Brazil | 25 | 999 | 6.31 | 0.001 | 0.001 |
| Kwale-Coast-Kenya | Sussex, UK | 35 | 999 | 18.80 | 0.001 | 0.001 |
| Laussane,<br>Switzerland | Nairobi-Kenya | 39 | 999 | 15.75 | 0.001 | 0.001 |
| Laussane,<br>Switzerland | S.J. Cariri , PB - Brazil | 19 | 999 | 3.47 | 0.001 | 0.001 |
| Laussane,<br>Switzerland | Sussex, UK | 29 | 999 | 6.73 | 0.001 | 0.001 |
| Nairobi-Kenya | S.J. Cariri , PB - Brazil | 30 | 999 | 6.61 | 0.001 | 0.001 |
| Nairobi-Kenya | Sussex, UK | 40 | 999 | 20.09 | 0.001 | 0.001 |
| S.J. Cariri , PB -<br>Brazil | Sussex, UK | 20 | 999 | 4.02 | 0.001 | 0.001 |

**Supplemental Table 3.** Pairwise comparisons of gut microbial communities by sampling location. There was a significant diference in gut microbial composition (Weighted UniFrac) between all sampling locations except the two locations in Brazil compared to each other and the four locations within Kenya compared to each other.

| Group 1 | Group 2 | Sample size | Permutations | pseudo-F | p-value | q-value |
| --- | --- | --- | --- | --- | --- | --- |
| Areia, PB - Brazil | Austin, Texas - US | 20 | 999 | 20.21 | 0.001 | 0.001 |
| Areia, PB - Brazil | Kakamega-Kenya | 23 | 999 | 8.25 | 0.001 | 0.001 |
| Areia, PB - Brazil | Kilifi-Coast-Kenya | 28 | 999 | 9.78 | 0.001 | 0.001 |
| Areia, PB - Brazil | Kwale-Coast-Kenya | 25 | 999 | 10.26 | 0.001 | 0.001 |
| Areia, PB - Brazil | Laussane, Switzerland | 19 | 999 | 19.21 | 0.001 | 0.001 |
| Areia, PB - Brazil | Nairobi-Kenya | 30 | 999 | 9.38 | 0.001 | 0.001 |
| Areia, PB - Brazil | S.J. Cariri , PB - Brazil | 10 | 999 | 1.21 | 0.273 | 0.315 |
| Areia, PB - Brazil | Sussex, UK | 20 | 999 | 10.20 | 0.002 | 0.002 |
| Austin, Texas - US | Kakamega-Kenya | 33 | 999 | 34.52 | 0.001 | 0.001 |
| Austin, Texas - US | Kilifi-Coast-Kenya | 38 | 999 | 39.31 | 0.001 | 0.001 |
| Austin, Texas - US | Kwale-Coast-Kenya | 35 | 999 | 41.50 | 0.001 | 0.001 |
| Austin, Texas - US | Laussane, Switzerland | 29 | 999 | 144.64 | 0.001 | 0.001 |
| Austin, Texas - US | Nairobi-Kenya | 40 | 999 | 38.46 | 0.001 | 0.001 |
| Austin, Texas - US | S.J. Cariri , PB - Brazil | 20 | 999 | 14.20 | 0.001 | 0.001 |
| Austin, Texas - US | Sussex, UK | 30 | 999 | 100.75 | 0.001 | 0.001 |
| Kakamega-Kenya | Kilifi-Coast-Kenya | 41 | 999 | 1.12 | 0.288 | 0.315 |
| Kakamega-Kenya | Kwale-Coast-Kenya | 38 | 999 | 1.71 | 0.051 | 0.061 |
| Kakamega-Kenya | Laussane, Switzerland | 32 | 999 | 25.78 | 0.001 | 0.001 |
| Kakamega-Kenya | Nairobi-Kenya | 43 | 999 | 1.14 | 0.289 | 0.315 |
| Kakamega-Kenya | S.J. Cariri , PB - Brazil | 23 | 999 | 8.97 | 0.001 | 0.001 |
| Kakamega-Kenya | Sussex, UK | 33 | 999 | 29.99 | 0.001 | 0.001 |
| Kilifi-Coast-Kenya | Kwale-Coast-Kenya | 43 | 999 | 0.74 | 0.772 | 0.794 |
| Kilifi-Coast-Kenya | Laussane, Switzerland | 37 | 999 | 30.43 | 0.001 | 0.001 |
| Kilifi-Coast-Kenya | Nairobi-Kenya | 48 | 999 | 0.64 | 0.883 | 0.883 |
| Kilifi-Coast-Kenya | S.J. Cariri , PB - Brazil | 28 | 999 | 10.63 | 0.001 | 0.001 |
| Kilifi-Coast-Kenya | Sussex, UK | 38 | 999 | 34.75 | 0.001 | 0.001 |
| Kwale-Coast-Kenya | Laussane, Switzerland | 34 | 999 | 31.92 | 0.001 | 0.001 |
| Kwale-Coast-Kenya | Nairobi-Kenya | 45 | 999 | 0.97 | 0.488 | 0.517 |
| Kwale-Coast-Kenya | S.J. Cariri , PB - Brazil | 25 | 999 | 11.12 | 0.001 | 0.001 |
| Kwale-Coast-Kenya | Sussex, UK | 35 | 999 | 36.43 | 0.001 | 0.001 |
| Laussane, Switzerland | Nairobi-Kenya | 39 | 999 | 29.37 | 0.001 | 0.001 |
| Laussane, Switzerland | S.J. Cariri , PB - Brazil | 19 | 999 | 36.17 | 0.001 | 0.001 |
| Laussane,<br>Switzerland | Sussex, UK | 29 | 999 | 8.08 | 0.001 | 0.001 |
| Nairobi-Kenya | S.J. Cariri , PB - Brazil | 30 | 999 | 10.26 | 0.001 | 0.001 |
| Nairobi-Kenya | Sussex, UK | 40 | 999 | 33.87 | 0.001 | 0.001 |
| S.J. Cariri , PB -<br>Brazil | Sussex, UK | 20 | 999 | 18.08 | 0.001 | 0.001 |

**Supplemental Table 4.** Differentially abundant microbiota by honey bee subspecies. Pairwise p-values (Kruskal-Wallis) comparing relative abundances of taxa identified as differentially abundant via ANCOM. Significant p-values are highlighted in green.

| Bacterial taxa | Pairwise *p*-values |  |  |
| --- | --- | --- | --- |
|  | European vs. African | European vs. Africanized | African vs. Africanized |
| *Frischella* | 3.8e-14 | 1 | < 2e-16 |
| *Gilliamella* | 0.0785 | 1 | 0.0046 |
| Orbaceae | 1.1e-14 | 3.7e-11 | 0.06 |
| Enterobacteriaceae | 0.0069 | 0.0021 | 0.2237 |
| *Pantoea* | 0.0013 | 0.4573 | - |
| Orbaceae | 3.9e-09 | 0.045 | - |
| Rhizobiaceae | 1.4e-13 | - | 0.00016 |
| Bombella | 0.00049 | 1 | 0.13864 |
| Bacteria | 7.4e-15 | - | 5.8e-05 |

**Supplemental Table 5.** Microbial species distinct to each honey bee subspecies. Microbial species were identified through an indicator species analysis (R, package indicspecies).

|  | Stat | p.value | Significance |
| --- | --- | --- | --- |
| **Group African** |  |  |  |
| Unclassified Bacteria | 0.732 | 0.0001 | *** |
| Proteobacteria; Alphaproteobacteria; Rhizobiales; Rhizobiaceae; Allorhizobium-Neorhizobium-Pararhizobium-Rhizobium | 0.621 | 0.0001 | *** |
| Proteobacteria; Alphaproteobacteria; Acetobacterales; Acetobacteraceae; Bombella | 0.406 | 0.0032 | ** |
| Proteobacteria; Gammaproteobacteria; Betaproteobacteriales; Burkholderiaceae | 0.276 | 0.0286 | * |
| **Group Africanized** |  |  |  |
| Proteobacteria; Gammaproteobacteria; Betaproteobacteriales; Neisseriaceae; Snodgrassella | 0.592 | 0.0002 | *** |
| Firmicutes; Bacilli; Lactobacillales; Lactobacillaceae; Lactobacillus | 0.588 | 0.0001 | *** |
| Actinobacteria; Actinobacteria; Bifidobacteriales; Bifidobacteriaceae; Bifidobacterium | 0.523 | 0.0002 | *** |
| Bacteroidetes; Bacteroidia; Flavobacteriales; Weeksellaceae; Apibacter | 0.306 | 0.0173 | * |
| **Group European** |  |  |  |
| Proteobacteria; Gammaproteobacteria; Orbales; Orbaceae | 0.515 | 0.0001 | *** |
| Proteobacteria; Gammaproteobacteria; Enterobacteriales; Enterobacteriaceae | 0.509 | 0.0001 | *** |
| Proteobacteria; Gammaproteobacteria; Enterobacteriales; Enterobacteriaceae; Pantoea | 0.308 | 0.0089 | ** |
| Proteobacteria; Gammaproteobacteria; Orbales; Orbaceae; Frischella | 0.273 | 0.0265 | * |
| **Group African+Africanized** |  |  |  |
| Proteobacteria; Gammaproteobacteria; Orbales; Orbaceae; uncultured gamma proteobacterium | 0.489 | 0.0001 | *** |
| **Group Africanized+European** |  |  |  |
| Proteobacteria; Gammaproteobacteria; Orbales; Orbaceae; Gilliamella | 0.546 | 0.0003 | *** |

