## Supplemental material: Scripts from R and Qiime2 for "Are you my mother?: When host genetics and gut microbiota tell different phylogenetic stories in the Africanized honey bee hybrid (*Apis mellifera scutellata* x sspp.)"

**Appendix 1**

**Script 1**

**QIIME script**

**Protocolo de analise *in silico* do artigo**

**“Africanized honey bees (*Apis mellifera scutellata* x sspp.) gut microbiota: is it close to African or European bees and what it implicates in bee health?”**

##PBS -N ondemand/sys/myjobs/default

#PBS -A PAS1331

#PBS -l walltime=48:00:00

#PBS -l nodes=1:ppn=10

#PBS -j oe

#PBS -m abe

\#

\# Move to the directory where the job was submitted

\#

\# Run script

\#

sh myscript.sh > my_results

\#

\# Now, copy data (or move) back once the simulation has completed

\# Run sequential job

\#

/usr/bin/time ./myscript.sh

\#

\# Now, copy data (or move) back once the simulation has completed

\#

cp message.out $SLURM_SUBMIT_DIR

\##

source activate qiime2-2020.2

#cd /users/PAS1331/kilmersoares/ondemand/data/sys/myjobs/projects/default/23

#Download and installing the sratoolki for ubuntu system

\# wget --output-document sratoolkit.tar.gz http://ftp-trace.ncbi.nlm.nih.gov/sra/sdk/current/sratoolkit.current-ubuntu64.tar.gz

#tar -vxzf sratoolkit.tar.gz

#export PATH=$PATH:/users/PAS1331/kilmersoares/ondemand/data/sys/myjobs/projects/default/23/sratoolkit.2.10.9-ubuntu64/bin

#which fastq-dump

#download the sra files

#STEP 1. Download a table of the metadata into a CSV file “SraRunInfo.csv”:

#From SRA web page :

#click on “Send to (top right corner)” Select “File” Select format “RunInfo” Click on “Create File”

#STEP 2. Read this CSV file “SraRunInfo.csv” into R:

#The SRA files are automatically download in the current working directory using the following R script:

#testing

#fastq-dump --stdout SRR10322593.1 | head -n 8

#mkdir readsSRA

#cd /users/PAS1331/kilmersoares/ondemand/data/sys/myjobs/projects/default/23/readsSRA

#save the file in a separeted directory

#cd /users/PAS1331/kilmersoares/ondemand/data/sys/myjobs/projects/default/23/(THE SEPARETED DIRECTORY CHOSEN)

#replace "texas.csv"for the name you Chosen

#R

#re

#Motta, 2018

#Read SRA file infos

#sri<-read.csv("texas.csv", stringsAsFactors=FALSE)

#head(sri)

#files<-basename(sri$download_path)

#for(i in 1:length(files)) download.file(sri$download_path[i], files[i])

\# Assure that all the files has been downloaded successfully

\# Remember, the R object files has been created in the previous code chunk

#stopifnot( all(file.exists(files)) )

#for(f in files) {

\# cmd = paste("fastq-dump --split-3", f)

#cat(cmd,"\n")#print the current command

#system(cmd) # invoke command

#Jones, 2018

#Read SRA file infos

#sri<-read.csv("uk.csv", stringsAsFactors=FALSE)

#head(sri)

#files<-basename(sri$download_path)

#for(i in 1:length(files)) download.file(sri$download_path[i], files[i])

\# Assure that all the files has been downloaded successfully

\# Remember, the R object files has been created in the previous code chunk

#stopifnot( all(file.exists(files)) )

#for(f in files) {

\# cmd = paste("fastq-dump --split-3", f)

\# cat(cmd,"\n")#print the current command

\# system(cmd) # invoke command

#}

#JKešnerová, 2019

#Read SRA file infos

#sri<-read.csv("Switzer.csv", stringsAsFactors=FALSE)

#head(sri)

#files<-basename(sri$download_path)

#for(i in 1:length(files)) download.file(sri$download_path[i], files[i])

\# Assure that all the files has been downloaded successfully

\# Remember, the R object files has been created in the previous code chunk

#stopifnot( all(file.exists(files)) )

#for(f in files) {

\# cmd = paste("fastq-dump --split-3", f)

\# cat(cmd,"\n")#print the current command

\# system(cmd) # invoke command

#}

#q()

cd /users/PAS1331/kilmersoares/ondemand/data/sys/myjobs/projects/default/23

#Importing data

#qiime tools import \

#--type 'SampleData[PairedEndSequencesWithQuality]' \

\# --input-path manifest_BR.csv \

\# --output-path demux_seqs_BR.qza \

\# --input-format PairedEndFastqManifestPhred33

\# qiime demux summarize \

\# --i-data demux_seqs_BR.qza \

\# --o-visualization demux_seqs_BR.qzv

\# qiime tools import \

\# --type 'SampleData[PairedEndSequencesWithQuality]' \

\# --input-path manifest_TX.csv \

\# --output-path demux_seqs_TX.qza \

\# --input-format PairedEndFastqManifestPhred33

\# qiime demux summarize \

\# --i-data demux_seqs_TX.qza \

\# --o-visualization demux_seqs_TX.qzv

\# qiime tools import \

\# --type 'SampleData[PairedEndSequencesWithQuality]' \

\# --input-path manifest_UK.csv \

\# --output-path demux_seqs_UK.qza \

\# --input-format PairedEndFastqManifestPhred33

\# qiime demux summarize \

\# --i-data demux_seqs_UK.qza \

\# --o-visualization demux_seqs_UK.qzv

\# qiime tools import \

\# --type 'SampleData[PairedEndSequencesWithQuality]' \

\# --input-path manifest_SW.csv \

\# --output-path demux_seqs_SW.qza \

\# --input-format PairedEndFastqManifestPhred33

\# qiime demux summarize \

\# --i-data demux_seqs_SW.qza \

\# --o-visualization demux_seqs_SW.qzv

\# qiime tools import \

\# --type 'SampleData[PairedEndSequencesWithQuality]' \

#--input-path manifest_KEcsv \

#--output-path demux_seqs_KE.qza \

#--input-format PairedEndFastqManifestPhred33

#qiime demux summarize \

\# --i-data demux_seqs_KE.qza \

\# --o-visualization demux_seqs_KE.qzv

#Sequence quality control and feature table

\# This method denoises paired-end sequences, dereplicates them, and filters chimeras.

#qiime dada2 denoise-paired \

\# --i-demultiplexed-seqs ./demux_seqs_BR.qza \

\# --p-trunc-len-f 250 \

\# --p-trunc-len-r 250 \

\# --o-table ./dada2_table_BR.qza \

\# --o-representative-sequences ./dada2_rep_set_BR.qza \

\# --o-denoising-stats ./dada2_stats_BR.qza

#qiime dada2 denoise-paired \

\# --i-demultiplexed-seqs ./demux_seqs_TX.qza \

\# --p-trunc-len-f 250 \

\# --p-trunc-len-r 250 \

\# --o-table ./dada2_table_TX.qza \

\# --o-representative-sequences ./dada2_rep_set_TX.qza \

\# --o-denoising-stats ./dada2_stats_TX.qza

#qiime dada2 denoise-paired \

\# --i-demultiplexed-seqs ./demux_seqs_UK.qza \

\# --p-trunc-len-f 220 \

\# --p-trunc-len-r 200 \

\# --o-table ./dada2_table_UK.qza \

\# --o-representative-sequences ./dada2_rep_set_UK.qza \

\# --o-denoising-stats ./dada2_stats_UK.qza

#qiime dada2 denoise-paired \

\# --i-demultiplexed-seqs ./demux_seqs_SW.qza \

\# --p-trunc-len-f 250 \

\# --p-trunc-len-r 250 \

\# --o-table ./dada2_table_SW.qza \

\# --o-representative-sequences ./dada2_rep_set_SW.qza \

\# --o-denoising-stats ./dada2_stats_SW.qza

#qiime dada2 denoise-paired \

\# --i-demultiplexed-seqs ./demux_seqs_KE.qza \

\# --p-trunc-len-f 250 \

\# --p-trunc-len-r 250 \

\# --o-table ./dada2_table_KE.qza \

\# --o-representative-sequences ./dada2_rep_set_KE.qza \

\# --o-denoising-stats ./dada2_stats_KE.qza

#METADATA DADA TABULATE####

qiime metadata tabulate \

--m-input-file metadata.tsv \

--o-visualization metadata.qzv

qiime metadata tabulate \

--m-input-file ./dada2_stats_BR.qza \

--o-visualization ./dada2_stats_BR.qzv

qiime feature-table summarize \

--i-table ./dada2_table_BR.qza \

--m-sample-metadata-file ./metadata_BR.tsv \

--o-visualization ./dada2_table_BR.qzv

qiime metadata tabulate \

--m-input-file ./dada2_stats_TX.qza \

--o-visualization ./dada2_stats_TX.qzv

qiime feature-table summarize \

--i-table ./dada2_table_TX.qza \

--m-sample-metadata-file ./metadata_TX.tsv \

--o-visualization ./dada2_table_TX.qzv

qiime metadata tabulate \

--m-input-file ./dada2_stats_UK.qza \

--o-visualization ./dada2_stats_UK.qzv

qiime feature-table summarize \

--i-table ./dada2_table_UK.qza \

--m-sample-metadata-file ./metadata_UK.tsv \

--o-visualization ./dada2_table_UK.qzv

qiime metadata tabulate \

--m-input-file ./dada2_stats_SW.qza \

--o-visualization ./dada2_stats_SW.qzv

qiime feature-table summarize \

--i-table ./dada2_table_SW.qza \

--m-sample-metadata-file ./metadata_SW.tsv \

--o-visualization ./dada2_table_SW.qzv

qiime metadata tabulate \

--m-input-file ./dada2_stats_KE.qza \

--o-visualization ./dada2_stats_KE.qzv

qiime feature-table summarize \

--i-table ./dada2_table_KE.qza \

--m-sample-metadata-file ./metadata_KE.tsv \

--o-visualization ./dada2_table_KE.qzv

#Alpha Rarefaction, Selecting a Rarefaction Depth

#Alpha Rarefaction, Selecting a Rarefaction Depth

\# Generate interactive alpha rarefaction curves by computing rarefactions between `min_depth` and `max_depth`. The number of intermediate depths to compute is controlled by the `steps` parameter, with n `iterations` being computed at each rarefaction depth. If sample metadata is provided, samples may be grouped based on distinct values within a metadata column

#We have chosen the values based on min and max the maximum sample total frequency of the feature_table dada2_table.qzv for each set of samples.

qiime diversity alpha-rarefaction \

--i-table ./dada2_table?.qza \

--m-metadata-file ./metadata_BR.tsv \

--o-visualization ./alpha_rarefaction_curves_BR.qzv \

--p-min-depth 1000 \

--p-max-depth	26837

qiime diversity alpha-rarefaction \

--i-table ./dada2_table_TX.qza \

--m-metadata-file ./metadata_TX.tsv \

--o-visualization ./alpha_rarefaction_curves_TX.qzv \

--p-min-depth 1000 \

--p-max-depth	26837

qiime diversity alpha-rarefaction \

--i-table ./dada2_table_UK.qza \

--m-metadata-file ./metadata_UK.tsv \

--o-visualization ./alpha_rarefaction_curves_UK.qzv \

--p-min-depth 5000 \

--p-max-depth	19000

qiime diversity alpha-rarefaction \

--i-table ./dada2_table_SW.qza \

--m-metadata-file ./metadata_SW.tsv \

--o-visualization ./alpha_rarefaction_curves_SW.qzv \

--p-min-depth 10000 \

--p-max-depth	18000

qiime diversity alpha-rarefaction \

--i-table ./dada2_table_KE.qza \

--m-metadata-file ./metadata_KE.tsv \

--o-visualization ./alpha_rarefaction_curves_KE.qzv \

--p-min-depth 1000 \

--p-max-depth	13000

#Generating a phylogenetic tree for diversity analysis

\# The following single command will produce two outputs: 1) phylogeny.qza is the Phylogeny[Rooted] and 2) placements.qza provides placement distributions for the fragments. Sequences which are not at least 75% similar by sequence identity to any record in the tree to insert into are not inserted into the tree.

\## We chose SEPP among the available phylogenetic insertion pipelines because of its scalable divide-and-conquer algorithm. recursively breaks down a problem into two or more sub-problems of the same or related type, until these become simple enough to be solved directly. The divide-and-conquer paradigm is often used to find an optimal solution of a problem.

#Generating a phylogenetic tree for diversity analysis BR

qiime fragment-insertion sepp \

--i-representative-sequences ./dada2_rep_set_BR.qza \

--i-reference-database sepp-refs-gg-13-8.qza \

--o-tree ./tree_BR.qza \

--o-placements ./tree_placements_BR.qza \

--p-threads 4

#creating core metrics results

\# Method, which rarefies a FeatureTable[Frequency] to a user-specified depth, computes several alpha and beta diversity metrics, and generates principle coordinates analysis (PCoA) plots using Emperor for each of the beta diversity metrics.

\# Generating a phylogenetic tree for diversity analysis BR

qiime diversity core-metrics-phylogenetic \

--i-table ./dada2_table_BR.qza \

--i-phylogeny ./tree_BR.qza \

--m-metadata-file ./metadata_BR.tsv \

--p-sampling-depth 1000 \

--output-dir ./core-metrics-results_BR

#Generating a phylogenetic tree for diversity analysis TX

qiime fragment-insertion sepp \

--i-representative-sequences ./dada2_rep_set_TX.qza \

--i-reference-database sepp-refs-gg-13-8.qza \

--o-tree ./tree_TX.qza \

--o-placements ./tree_placements_TX.qza \

--p-threads 4

qiime diversity core-metrics-phylogenetic \

--i-table ./dada2_table_TX.qza \

--i-phylogeny ./tree_TX.qza \

--m-metadata-file ./metadata_TX.tsv \

--p-sampling-depth 1600 \

--output-dir ./core-metrics-results_TX

#Generating a phylogenetic tree for diversity analysis UK

qiime fragment-insertion sepp \

--i-representative-sequences ./dada2_rep_set_UK.qza \

--i-reference-database sepp-refs-gg-13-8.qza \

--o-tree ./tree_UK.qza \

--o-placements ./tree_placements_UK.qza \

--p-threads 4

qiime diversity core-metrics-phylogenetic \

--i-table ./dada2_table_UK.qza \

--i-phylogeny ./tree_UK.qza \

--m-metadata-file ./metadata_UK.tsv \

--p-sampling-depth 5000 \

--output-dir ./core-metrics-results_UK

#Generating a phylogenetic tree for diversity analysis SW

qiime fragment-insertion sepp \

--i-representative-sequences ./dada2_rep_set_SW.qza \

--i-reference-database sepp-refs-gg-13-8.qza \

--o-tree ./tree_SW.qza \

--o-placements ./tree_placements_SW.qza \

--p-threads 4

qiime diversity core-metrics-phylogenetic \

--i-table ./dada2_table_SW.qza \

--i-phylogeny ./tree_SW.qza \

--m-metadata-file ./metadata_SW.tsv \

--p-sampling-depth 10000 \

--output-dir ./core-metrics-results_SW

#Generating a phylogenetic tree for diversity analysis KE

qiime fragment-insertion sepp \

--i-representative-sequences ./dada2_rep_set_KE.qza \

--i-reference-database sepp-refs-gg-13-8.qza \

--o-tree ./tree_KE.qza \

--o-placements ./tree_placements_KE.qza \

--p-threads 4

qiime diversity core-metrics-phylogenetic \

--i-table ./dada2_table_KE.qza \

--i-phylogeny ./tree_KE.qza \

--m-metadata-file ./metadata_KE.tsv \

--p-sampling-depth 1200 \

--output-dir ./core-metrics-results_KE

#Diversity analysis

#Alpha diversity

##We used the alpha group significance plugin to test for differences in alpha diversity

qiime diversity alpha-group-significance \

--i-alpha-diversity ./core-metrics-results_BR/faith_pd_vector.qza \

--m-metadata-file ./metadata_BR.tsv \

--o-visualization ./core-metrics-results_BR/faiths_pd_statistics_BR.qzv

qiime diversity alpha-group-significance \

--i-alpha-diversity ./core-metrics-results_BR/evenness_vector.qza \

--m-metadata-file ./metadata_BR.tsv \

--o-visualization ./core-metrics-results_BR/evenness_statistics_BR.qzv

qiime diversity alpha-group-significance \

--i-alpha-diversity ./core-metrics-results_BR/shannon_vector.qza \

--m-metadata-file ./metadata_BR.tsv \

--o-visualization ./core-metrics-results_BR/shannon_statistics_BR.qzv

qiime diversity alpha-group-significance \

--i-alpha-diversity ./core-metrics-results_BR/observed_otus_vector.qza \

--m-metadata-file ./metadata_BR.tsv \

--o-visualization ./core-metrics-results_BR/observed_otus_vector_BR.qzv

qiime diversity alpha-group-significance \

--i-alpha-diversity ./core-metrics-results_TX/faith_pd_vector.qza \

--m-metadata-file ./metadata_TX.tsv \

--o-visualization ./core-metrics-results_TX/faiths_pd_statistics_TX.qzv

qiime diversity alpha-group-significance \

--i-alpha-diversity ./core-metrics-results_TX/evenness_vector.qza \

--m-metadata-file ./metadata_TX.tsv \

--o-visualization ./core-metrics-results_TX/evenness_statistics_TX.qzv

qiime diversity alpha-group-significance \

--i-alpha-diversity ./core-metrics-results_TX/shannon_vector.qza \

--m-metadata-file ./metadata_TX.tsv \

--o-visualization ./core-metrics-results_TX/shannon_statistics_TX.qzv

qiime diversity alpha-group-significance \

--i-alpha-diversity ./core-metrics-results_TX/observed_otus_vector.qza \

--m-metadata-file ./metadata_TX.tsv \

--o-visualization ./core-metrics-results_TX/observed_otus_vector_TX.qzv

#Alpha diversity

qiime diversity alpha-group-significance \

--i-alpha-diversity ./core-metrics-results_UK/faith_pd_vector.qza \

--m-metadata-file ./metadata_UK.tsv \

--o-visualization ./core-metrics-results_UK/faiths_pd_statistics_UK.qzv

qiime diversity alpha-group-significance \

--i-alpha-diversity ./core-metrics-results_UK/evenness_vector.qza \

--m-metadata-file ./metadata_UK.tsv \

--o-visualization ./core-metrics-results_UK/evenness_statistics_UK.qzv

qiime diversity alpha-group-significance \

--i-alpha-diversity ./core-metrics-results_UK/shannon_vector.qza \

--m-metadata-file ./metadata_UK.tsv \

--o-visualization ./core-metrics-results_UK/shannon_statistics_UK.qzv

qiime diversity alpha-group-significance \

--i-alpha-diversity ./core-metrics-results_UK/observed_otus_vector.qza \

--m-metadata-file ./metadata_UK.tsv \

--o-visualization ./core-metrics-results_UK/observed_otus_vector_UK.qzv

qiime diversity alpha-group-significance \

--i-alpha-diversity ./core-metrics-results_SW/faith_pd_vector.qza \

--m-metadata-file ./metadata_SW.tsv \

--o-visualization ./core-metrics-results_SW/faiths_pd_statistics_SW.qzv

qiime diversity alpha-group-significance \

--i-alpha-diversity ./core-metrics-results_SW/evenness_vector.qza \

--m-metadata-file ./metadata_SW.tsv \

--o-visualization ./core-metrics-results_SW/evenness_statistics_SW.qzv

qiime diversity alpha-group-significance \

--i-alpha-diversity ./core-metrics-results_SW/shannon_vector.qza \

--m-metadata-file ./metadata_SW.tsv \

--o-visualization ./core-metrics-results_SW/shannon_statistics_SW.qzv

qiime diversity alpha-group-significance \

--i-alpha-diversity ./core-metrics-results_SW/observed_otus_vector.qza \

--m-metadata-file ./metadata_SW.tsv \

--o-visualization ./core-metrics-results_SW/observed_otus_vector_SW.qzv

qiime diversity alpha-group-significance \

--i-alpha-diversity ./core-metrics-results_KE/faith_pd_vector.qza \

--m-metadata-file ./metadata_KE.tsv \

--o-visualization ./core-metrics-results_KE/faiths_pd_statistics_KE.qzv

qiime diversity alpha-group-significance \

--i-alpha-diversity ./core-metrics-results_KE/evenness_vector.qza \

--m-metadata-file ./metadata_KE.tsv \

--o-visualization ./core-metrics-results_KE/evenness_statistics_KE.qzv

qiime diversity alpha-group-significance \

--i-alpha-diversity ./core-metrics-results_KE/shannon_vector.qza \

--m-metadata-file ./metadata_KE.tsv \

--o-visualization ./core-metrics-results_KE/shannon_statistics_KE.qzv

qiime diversity alpha-group-significance \

--i-alpha-diversity ./core-metrics-results_KE/observed_otus_vector.qza \

--m-metadata-file ./metadata_KE.tsv \

--o-visualization ./core-metrics-results_KE/observed_otus_vector_KE.qzv

#MERGING for overall analyses

##We merged all tables for downstream analyses

qiime feature-table merge \

--i-tables dada2_table_BR.qza \

--i-tables dada2_table_TX.qza \

--i-tables dada2_table_UK.qza \

--i-tables dada2_table_SW.qza \

--i-tables dada2_table_KE.qza \

--o-merged-table dada2_table.qza

qiime feature-table merge-seqs \

--i-data dada2_rep_set_BR.qza \

--i-data dada2_rep_set_TX.qza \

--i-data dada2_rep_set_UK.qza \

--i-data dada2_rep_set_SW.qza \

--i-data dada2_rep_set_KE.qza \

--o-merged-data dada2_rep_set.qza

qiime feature-table summarize \

--i-table ./dada2_table.qza \

--m-sample-metadata-file ./metadata.tsv \

--o-visualization ./dada2_table.qzv

#Generating a phylogenetic tree for diversity analysis

qiime fragment-insertion sepp \

--i-representative-sequences ./dada2_rep_set.qza \

--i-reference-database sepp-refs-gg-13-8.qza \

--o-tree ./tree.qza \

--o-placements ./tree_placements.qza \

--p-threads 4 # update to a higher number if you can

##I have chosen I have chosen 1000 and 26959 based on min and max the maximum sample total frequency of the feature_table dada2_table.qzv. The minimum rarefaction depth. The maximum rarefaction depth must be greater than min-depth.

Range(1, None) min-depth.

qiime diversity alpha-rarefaction \

--i-table ./dada2_table.qza \

--m-metadata-file ./metadata.tsv \

--o-visualization ./alpha_rarefaction_curves.qzv \

--p-min-depth 1000 \

--p-max-depth 26959

#Diversity analysis

#I have chosen p-sampling-depth 1000 based on the min Feature Count on dada2_table.qzv

qiime diversity core-metrics-phylogenetic \

--i-table ./dada2_table.qza \

--i-phylogeny ./tree.qza \

--m-metadata-file ./metadata.tsv \

--p-sampling-depth 1000 \

--output-dir ./core-metrics-results

#Alpha Rarefaction group significance (after core metrics).

\## We will test for associations between categories in the sample metadata file and alpha diversity data.

qiime diversity alpha-group-significance \

--i-alpha-diversity core-metrics-results/shannon_vector.qza \

--m-metadata-file metadata.tsv \

--o-visualization core-metrics-results/shannon_vector-group-significance.qzv

#Alpha diversity

qiime diversity alpha-group-significance \

--i-alpha-diversity ./core-metrics-results/faith_pd_vector.qza \

--m-metadata-file ./metadata.tsv \

--o-visualization ./core-metrics-results/faiths_pd_statistics.qzv

qiime diversity alpha-group-significance \

--i-alpha-diversity ./core-metrics-results/evenness_vector.qza \

--m-metadata-file ./metadata.tsv \

--o-visualization ./core-metrics-results/evenness_statistics.qzv

qiime diversity alpha-group-significance \

--i-alpha-diversity ./core-metrics-results/shannon_vector.qza \

--m-metadata-file ./metadata.tsv \

--o-visualization ./core-metrics-results/shannon_statistics.qzv

qiime diversity alpha-group-significance \

--i-alpha-diversity ./core-metrics-results/observed_otus_vector.qza \

--m-metadata-file ./metadata.tsv \

--o-visualization ./core-metrics-results/observed_otus_statistic.qzv

#Beta diversity OVERALL ANALISES

\# Determine whether groups of samples are significantly different from one another using a permutation-based statistical test.

#For biome

qiime diversity beta-group-significance \

--i-distance-matrix core-metrics-results/unweighted_unifrac_distance_matrix.qza \

--m-metadata-file metadata.tsv \

--m-metadata-column biome \

--p-pairwise \

--o-visualization core-metrics-results/unweighted-unifrac-biome-significance.qzv

qiime diversity beta-group-significance \

--i-distance-matrix core-metrics-results/weighted_unifrac_distance_matrix.qza \

--m-metadata-file metadata.tsv \

--m-metadata-column biome \

--p-pairwise \

--o-visualization core-metrics-results/weighted-unifrac-biome-significance.qzv

qiime diversity beta-group-significance \

--i-distance-matrix core-metrics-results/bray_curtis_distance_matrix.qza \

--m-metadata-file metadata.tsv \

--m-metadata-column biome \

--p-pairwise \

--o-visualization core-metrics-results/bray_curtis-biome-significance.qzv

qiime diversity beta-group-significance \

--i-distance-matrix core-metrics-results/jaccard_distance_matrix.qza \

--m-metadata-file metadata.tsv \

--m-metadata-column biome \

--p-pairwise \

--o-visualization core-metrics-results/jaccard-biome-significance_pairwise.qzv

#For Subspecie

qiime diversity beta-group-significance \

--i-distance-matrix core-metrics-results/unweighted_unifrac_distance_matrix.qza \

--m-metadata-file metadata.tsv \

--m-metadata-column species \

--o-visualization core-metrics-results/unweighted-unifrac-species-significance.qzv \

--p-pairwise

qiime diversity beta-group-significance \

--i-distance-matrix core-metrics-results/weighted_unifrac_distance_matrix.qza \

--m-metadata-file metadata.tsv \

--m-metadata-column species \

--p-pairwise \

--o-visualization core-metrics-results/weighted-unifrac-species-significance.qzv \

qiime diversity beta-group-significance \

--i-distance-matrix core-metrics-results/bray_curtis_distance_matrix.qza \

--m-metadata-file metadata.tsv \

--m-metadata-column species \

--p-pairwise \

--o-visualization core-metrics-results/bray_curtis-species-significance.qzvI

qiime diversity beta-group-significance \

--i-distance-matrix core-metrics-results/jaccard_distance_matrix.qza \

--m-metadata-file metadata.tsv \

--m-metadata-column species \

--p-pairwise \

--o-visualization core-metrics-results/jaccard-species-significance_pairwise.qzv

#Adonis (PERMANOVA)

\## It is a non-parametric multivariate statistical permutation test. PERMANOVA is used to compare groups of objects and test the null hypothesis that the centroids and dispersion of the groups as defined by measure space are equivalent for all groups. We used Adonis to look at a multivariate model. to see the effect of more than one grouping variable.

qiime diversity adonis \

--i-distance-matrix core-metrics-results/jaccard_distance_matrix.qza \

--m-metadata-file metadata.tsv \

--o-visualization core-metrics-results/jaccard_adonis.qzv \

--p-formula species

qiime diversity adonis \

--i-distance-matrix core-metrics-results/bray_curtis_distance_matrix.qza \

--m-metadata-file metadata.tsv \

--o-visualization core-metrics-results/bray_curtis_adonis.qzv \

--p-formula species

qiime diversity adonis \

--i-distance-matrix core-metrics-results/unweighted_unifrac_distance_matrix.qza \

--m-metadata-file metadata.tsv \

--o-visualization core-metrics-results/unweighted_unifrac_distance_adonis.qzv \

--p-formula species

qiime diversity adonis \

--i-distance-matrix core-metrics-results/weighted_unifrac_distance_matrix.qza \

--m-metadata-file metadata.tsv \

--o-visualization core-metrics-results/weighted_unifrac_distance_adonis.qzv \

--p-formula species

#Taxonomic classification

#alinhar e classificar

#Plugin for taxonomic classification of sequences. Contains multiple methods for sequence classification, including methods to train and employ scikit-learn classifiers for sequence classification.

#The naive Bayes algorithm for multinomially distributed data, and is one of the two classic naive Bayes variants used in text classification (where the data are typically represented as word vector counts, although tf-idf vectors are also known to work well in practice).

#We use silva-132-99-nb-classifier.qza

qiime feature-classifier classify-sklearn \

--i-reads ./dada2_rep_set.qza \

--i-classifier ./silva-132-99-nb-classifier.qza \

--o-classification ./taxonomy.qza

qiime metadata tabulate \

--m-input-file ./taxonomy.qza \

--o-visualization ./taxonomy.qzv

qiime feature-table tabulate-seqs \

--i-data ./dada2_rep_set.qza \

--o-visualization ./dada2_rep_set.qzv

#Filter the table of ASVs . Filtering can be applied to retain only specific taxa.

#Mitochondria

qiime taxa filter-table \

--i-table dada2_table.qza \

--i-taxonomy taxonomy.qza \

--p-exclude	Mitochondria \

--o-filtered-table table-no-mitochondria.qza

#Chloroplast

qiime taxa filter-table \

--i-table table-no-mitochondria.qza \

--i-taxonomy taxonomy.qza \

--p-exclude	Chloroplast \

--o-filtered-table table-no-chloroplast.qza

#Unassigned

qiime taxa filter-table \

--i-table table-no-chloroplast.qza \

--i-taxonomy taxonomy.qza \

--p-exclude	Unassigned \

--o-filtered-table table-no-Unassigned.qza

#Eukaryota

qiime taxa filter-table \

--i-table table-no-Unassigned.qza \

--i-taxonomy taxonomy.qza \

--p-exclude	Eukaryota \

--o-filtered-table table-final.qza

\# reattribute taxonomy

qiime feature-table filter-seqs \

--i-data dada2_rep_set.qza \

--i-table table-final.qza \

--o-filtered-data seqs-final.qza

#Taxonomy barchart (5000 based on the min Feature Count on dada2_table.qzv

qiime feature-table filter-samples \

--i-table ./table-final.qza \

--p-min-frequency 1000 \

--o-filtered-table ./table_2kfiltered.qza

qiime taxa barplot \

--i-table ./table_2kfiltered.qza \

--i-taxonomy ./taxonomy.qza \

--m-metadata-file ./metadata.tsv \

--o-visualization ./taxa_barplotfiltered.qzv

#Generates the table with the number of asvs per sample And the taxa in it and with the first table. We use the taxa table collapsed to the ASV level

\# table dada2 with taxonomy

qiime taxa collapse \

--i-table table-final.qza \

--i-taxonomy taxonomy.qza \

--p-level 7 \

--o-collapsed-table table-l7.qza

#Generates the table with the number of ASVs per sample

qiime tools export \

--input-path table-l7.qza \

--output-path table_exported-feature-table_L7

cd table_exported-feature-table_L7

biom convert --to-tsv -i feature-table.biom -o bee_sub_table_L7.tsv

cd /users/PAS1331/kilmersoares/ondemand/data/sys/myjobs/projects/default/23

#Differential abundance with ANCOM Analysis of composition of microbiomes. Filter features from table based on frequency and/or metadata. Any samples with a frequency of zero after feature filtering will also be removed.

\## p-min-frequency: The minimum total frequency that a feature must have o be retained; --p-max-frequency: The maximum total frequency that a feature can have to be retained. If no value is provided this will default to infinity (i.e., no maximum frequency filter will be applied).

qiime feature-table filter-features \

--i-table table_2kfiltered.qza \

--p-min-frequency 50 \

--p-min-samples 5 \

--o-filtered-table table_2k_abund.qza

\# table with taxonomy

qiime taxa collapse \

--i-table table_2k_abund.qza \

--i-taxonomy taxonomy.qza \

--p-level 6 \

--o-collapsed-table table-l62.qza

#Differential abundance with ANCOM

\## ANCOM accounts for the underlying structure in the data and can be used for comparing the composition of microbiomes in two or more populations.

\# First, incrementing all counts in table by pseudocount.

qiime composition add-pseudocount \

--i-table table-l62.qza \

--o-composition-table comp-table-l62.qza

#Ancon biome

qiime composition ancom \

--i-table comp-table-l62.qza \

--m-metadata-file metadata.tsv \

--m-metadata-column biome \

--o-visualization ancom-subject_biome.qzv

#Ancon species

qiime composition ancom \

--i-table comp-table-l62.qza \

--m-metadata-file metadata.tsv \

--m-metadata-column species \

--o-visualization ancom-subject_species.qzv

#Core features

\# Identify "core" features, which are features observed in a user-defined fraction of the samples.

#We considerered ASVs present in at least 90% of the samples

qiime feature-table core-features \

--i-table table-l6.qza \

--o-visualization core-features

#Machine-learning classifiers for predicting sample characteristics

\# Predicts a categorical sample metadata column using a supervised learning classifier. Uses nested stratified k-fold cross validation for automated hyperparameter optimization and sample prediction. Outputs predicted values for each input sample, and relative importance of each feature for model accuracy.

qiime sample-classifier classify-samples \

--i-table ./table-l6.qza \

--m-metadata-file ./metadata.tsv \

--m-metadata-column species \

--p-random-state 666 \

--p-n-jobs 1 \

--output-dir ./sample-classifier-results/

#Heat map

\## Generate a heatmap of important features. Features are filtered based on importance scores; samples are optionally grouped by metadata; and a heatmap is generated that displays (normalized) feature abundances per sample.

qiime sample-classifier heatmap \

--i-table ./table-l6.qza \

--i-importance ./sample-classifier-results/feature_importance.qza \

--m-sample-metadata-file metadata.tsv \

--m-sample-metadata-column species \

--p-group-samples \

--p-feature-count 100 \

--o-heatmap ./sample-classifier-results/heatmap100.qzv \

--o-filtered-table ./sample-classifier-results/filtered-table100.qza

qiime sample-classifier heatmap \

--i-table ./table-l6.qza \

--i-importance ./sample-classifier-results/feature_importance.qza \

--m-sample-metadata-file metadata.tsv \

--m-sample-metadata-column species \

--p-group-samples \

--p-feature-count 50 \

--o-heatmap ./sample-classifier-results/heatmap50.qzv \

--o-filtered-table ./sample-classifier-results/filtered-table50.qza

qiime sample-classifier heatmap \

--i-table ./table-l6.qza \

--i-importance ./sample-classifier-results/feature_importance.qza \

--m-sample-metadata-file metadata.tsv \

--m-sample-metadata-column species \

--p-group-samples \

--p-feature-count 10 \

--o-heatmap ./sample-classifier-results/heatmap10.qzv \

--o-filtered-table ./sample-classifier-results/filtered-table10.qza

qiime sample-classifier heatmap \

--i-table ./table-l6.qza \

--i-importance ./sample-classifier-results/feature_importance.qza \

--m-sample-metadata-file metadata.tsv \

--m-sample-metadata-column species \

--p-group-samples \

--p-feature-count 20 \

--o-heatmap ./sample-classifier-results/heatmap20.qzv \

--o-filtered-table ./sample-classifier-results/filtered-table20.qza

qiime sample-classifier heatmap \

--i-table ./table-l6.qza \

--i-importance ./sample-classifier-results/feature_importance.qza \

--m-sample-metadata-file metadata.tsv \

--m-sample-metadata-column species \

--p-group-samples \

--p-feature-count 15 \

--o-heatmap ./sample-classifier-results/heatmap15.qzv \

--o-filtered-table ./sample-classifier-results/filtered-table15.qza

**Script 2**

**R studio Post hoc analysis**

**Protocolo de analise *in silico* do artigo**

**“Africanized honey bees (*Apis mellifera scutellata* x sspp.) gut microbiota: is it close to African or European bees and what it implicates in bee health?”**

#importdata

library(readr)

library(readr)

data_sub <- read_csv("~/DOUTORADO/PRINT/ESTUDY/COmputationalanalises/Subspecies/Sub_R/data_sub.csv")

View(data_sub)

#Tukey test

library(laercio)

require(laercio)

#Subspecies

#diversity

par(mfrow = c(1,2), oma = c(2,1,1,1))

\# shannon

shapiro.test(data_sub$shannon) #not signifcant, data are normal

#modelo

#anova <- aov(shannon ~ Abrev, data=data_sub)

#summary(anova)

#tukey_ctrl <- LTukey(anova, which="Abrev", conf.level=0.95)

#kruskal.test

kruskal.test(shannon ~ Abrev, data=data_sub)

pairwise.wilcox.test(data_sub $ shannon,data_sub $Abrev, p.adj = "bonferroni", exact=FALSE)

#PLOT

boxplot(shannon ~ Abrev, data=data_sub, cex = 1, main=" Shannon's Index",xlab="Subspecies", ylab="Shannon's diversity index", col = c("#00CC00","#ffff00","#ff3300"))

#faith

shapiro.test(data_sub$faith) #not signifcant, data are normal

#modelo

#anova <- aov(faith ~ Abrev, data=data_sub)

#summary(anova)

#tukey_ctrl <- LTukey(anova, which="Abrev", conf.level=0.95)

#kruskal.test

kruskal.test(faith ~ Abrev, data=data_sub)

pairwise.wilcox.test(data_sub $ faith,data_sub $Abrev, p.adj = "bonferroni")

#PLOT

boxplot(faith ~ Abrev, data=data_sub, cex = 1, main= "Faith", xlab="Subspecies", ylab= "Faiths's diversity index",col = c("#00CC00","#ffff00","#ff3300"))

\# pielou

shapiro.test(data_sub$pielou) #not signifcant, data are normal

#modelo

#anova <- aov(pielou ~ Abrev, data=data_sub)

#summary(anova)

#tukey_ctrl <- LTukey(anova, which="Abrev",conf.level=0.95)

#kruskal.test

kruskal.test(pielou ~ Abrev, data=data_sub)

pairwise.wilcox.test(data_sub $ pielou,data_sub $Abrev, p.adj = "bonferroni", exact=FALSE)

#PLOT

boxplot(pielou ~ Abrev, data=data_sub, cex = 1, main="Eveness Pielou",xlab="Subspecies", ylab="Pielou's diversity index", col = c("#00CC00","#ffff00","#ff3300"))

#observed_otus

shapiro.test(data_sub$observed_otus) #not signifcant, data are normal

##modelo

#anova <- aov(observed_otus ~ Abrev, data=data_sub)

#summary(anova)

#tukey_ctrl <- LTukey(anova, which="Abrev", conf.level=0.95)

#kruskal.test

kruskal.test(observed_otus ~ Abrev, data=data_sub)

pairwise.wilcox.test(data_sub $observed_otus, data_sub $Abrev, p.adj = "bonferroni", exact=FALSE)

#PLOT

boxplot(observed_otus ~ Abrev, data=data_sub, cex = 1, main=" Observed ASVs",xlab="Subspecies", ylab="Observed ASVs", col = c("#00CC00","#ffff00","#ff3300"))

#relative abundances

#CORE

par(mfrow = c(1,3), oma = c(2,1,1,1))

#Lactobacillus

shapiro.test(data_sub$Lactobacillus) #not signifcant, data are normal

#modelo

#anova <- aov(Lactobacillus ~ Abrev, data=data_sub)

#summary(anova)

#tukey_ctrl <- LTukey(anova, which="Abrev", conf.level=0.95)

#kruskal.test

kruskal.test(species ~ Lactobacillus, data=data_sub)

pairwise.wilcox.test(data_sub $ Lactobacillus, data_sub $Abrev, p.adj = "bonferroni", exact=FALSE)

#PLOT

boxplot(Lactobacillus ~ Abrev, data=data_sub, cex = 1, main=" Lactobacillus",xlab="Subspecies", ylab="Relative abundance (%)", col = c("#00CC00","#ffff00","#ff3300"))

\# Snodgrassella

shapiro.test(data_sub$Snodgrassella) #not signifcant, data are normal

#modelo

#anova <- aov(Snodgrassella ~ Abrev, data=data_sub)

#summary(anova)

#tukey_ctrl <- LTukey(anova, which="Abrev", conf.level=0.95)

#kruskal.test

kruskal.test(species ~ Snodgrassella, data=data_sub)

pairwise.wilcox.test(data_sub $ Snodgrassella, data_sub $Abrev, p.adj = "bonferroni", exact=FALSE)

#PLOT

boxplot(Snodgrassella ~ Abrev, data=data_sub, cex = 1, main=" Snodgrassella",xlab="Subspecies", ylab="Relative abundance (%)", col = c("#00CC00","#ffff00","#ff3300"))

\# Gilliamella ####CORE AND ANCOM

shapiro.test(data_sub$Gilliamella) #not signifcant, data are not normal

#modelo

#anova <- aov(Gilliamella ~ Abrev, data=data_sub)

#summary(anova)

#tukey_ctrl <- LTukey(anova, which="Abrev", conf.level=0.95)

#kruskal.test

kruskal.test(species ~ Gilliamella, data=data_sub)

pairwise.wilcox.test(data_sub $ Gilliamella, data_sub $Abrev, p.adj = "bonferroni", exact=FALSE)

#PLOT

boxplot(Gilliamella ~ Abrev, data=data_sub, cex = 1, main=" Gilliamella",xlab="Subspecies", ylab="Relative abundance (%)", col = c("#00CC00","#ffff00","#ff3300"))

#ANCOM

par(mfrow = c(1,1), oma = c(2,1,1,1))

\#

\# Frischella

shapiro.test(data_sub$Frischella) #not signifcant, data are normal

#modelo

#anova <- aov(Frischella ~ Abrev, data=data_sub)

#summary(anova)

#tukey_ctrl <- LTukey(anova, which="Abrev", conf.level=0.95)

#kruskal.test

kruskal.test(species ~ Frischella, data=data_sub)

pairwise.wilcox.test(data_sub $ Frischella, data_sub $Abrev, p.adj = "bonferroni", exact=FALSE)

#PLOT

boxplot(Frischella ~ Abrev, data=data_sub, cex = 1, main=" Frischella",xlab="Subspecies", ylab="Relative abundance (%)", col = c("#00CC00","#ffff00","#ff3300"))

\# Uncultured gamma proteobacterium

shapiro.test(data_sub$Uncultured_gamma_proteobacterium) #not signifcant, data are normal

#modelo

#anova <- aov(Uncultured_gamma_proteobacterium ~ Abrev, data=data_sub)

#summary(anova)

#tukey_ctrl <- LTukey(anova, which="Abrev", conf.level=0.95)

#kruskal.test

kruskal.test(species ~ Uncultured_gamma_proteobacterium , data=data_sub)

pairwise.wilcox.test(data_sub $ Uncultured_gamma_proteobacterium , data_sub $Abrev, p.adj = "bonferroni", exact=FALSE)

#PLOT

boxplot(Uncultured_gamma_proteobacterium ~ Abrev, data=data_sub, cex = 1, main="Uncultured_gamma_proteobacterium ", xlab="Subspecies", ylab="Relative abundance (%)", col = c("#00CC00","#ffff00","#ff3300"))

\# Enterobacteriaceae

shapiro.test(data_sub$Enterobacteriaceae) #not signifcant, data are normal

#modelo

#anova <- aov(Enterobacteriaceae ~ Abrev, data=data_sub)

#summary(anova)

#tukey_ctrl <- LTukey(anova, which="Abrev", conf.level=0.95)

#kruskal.test

kruskal.test(species ~ Enterobacteriaceae, data=data_sub)

pairwise.wilcox.test(data_sub $ Enterobacteriaceae, data_sub $Abrev, p.adj = "bonferroni", exact=FALSE)

#PLOT

boxplot(Enterobacteriaceae ~ Abrev, data=data_sub, cex = 1, main=" Enterobacteriaceae",xlab="Subspecies", ylab="Relative abundance (%)", col = c("#00CC00","#ffff00","#ff3300"))

\# Rhizobiaceae

shapiro.test(data_sub$Rhizobiaceae) #not signifcant, data are normal

#modelo

#anova <- aov(Rhizobiaceae ~ Abrev, data=data_sub)

#summary(anova)

#tukey_ctrl <- LTukey(anova, which="Abrev", conf.level=0.95)

#kruskal.test

kruskal.test(species ~ Rhizobiaceae, data=data_sub)

pairwise.wilcox.test(data_sub $ Rhizobiaceae,data_sub $Abrev, p.adj = "bonferroni", exact=FALSE)

#PLOT

boxplot(Rhizobiaceae ~ Abrev, data=data_sub, cex = 1, main=" Rhizobiaceae",xlab="Subspecies", ylab="Relative abundance (%)", col = c("#00CC00","#ffff00","#ff3300"))

\# Pantoea

shapiro.test(data_sub$Pantoea) #not signifcant, data are normal

#modelo

anova <- aov(Pantoea ~ Abrev, data=data_sub)

summary(anova)

tukey_ctrl <- LTukey(anova, which="Abrev", conf.level=0.95)

#kruskal.test

kruskal.test(species ~ Pantoea, data=data_sub)

pairwise.wilcox.test(data_sub $ Pantoea, data_sub $Abrev, p.adj = "bonferroni", exact=FALSE)

#PLOT

boxplot(Pantoea ~ Abrev, data=data_sub, cex = 1, main=" Pantoea ",xlab="Subspecies", ylab="Relative abundance (%)", col = c("#00CC00","#ffff00","#ff3300"))

\# Bombella

shapiro.test(data_sub$Bombella) #not signifcant, data are normal

#modelo

anova <- aov(Bombella ~ Abrev, data=data_sub)

summary(anova)

tukey_ctrl <- LTukey(anova, which="Abrev", conf.level=0.95)

#kruskal.test

kruskal.test(species ~ Bombella, data=data_sub)

pairwise.wilcox.test(data_sub $ Bombella,data_sub $Abrev, p.adj = "bonferroni", exact=FALSE)

#PLOT

boxplot(Bombella ~ Abrev, data=data_sub, cex = 1, main=" Bombella",xlab="Subspecies", ylab="Relative abundance (%)", col = c("#00CC00","#ffff00","#ff3300"))

Gilliamella ####CORE AND ANCOM

shapiro.test(data_sub$Gilliamella) #not signifcant, data are not normal

modelo

anova <- aov(Gilliamella ~ Abrev, data=data_sub)

summary(anova)

tukey_ctrl <- LTukey(anova, which="Abrev", conf.level=0.95)

#kruskal.test

kruskal.test(species ~ Gilliamella, data=data_sub)

pairwise.wilcox.test(data_sub $ Gilliamella, data_sub $Abrev, p.adj = "bonferroni", exact=FALSE)

#PLOT

boxplot(Gilliamella ~ Abrev, data=data_sub, cex = 1, main=" Gilliamella",xlab="Subspecies", ylab="Relative abundance (%)", col = c("#00CC00","#ffff00","#ff3300"))

\# Bacteria

shapiro.test(data_sub$ Bacteria) #not signifcant, data are normal

#modelo

anova <- aov(Bacteria ~ Abrev, data=data_sub)

summary(anova)

tukey <- LTukey(anova, which="Abrev", conf.level=0.95)

#kruskal.test

kruskal.test(species ~ Bacteria, data=data_sub)

pairwise.wilcox.test(data_sub $ Bacteria, data_sub $Abrev, p.adj = "bonferroni", exact=FALSE)

#PLOT

boxplot(Bacteria ~ Abrev, data=data_sub, cex = 1, main=" Bacteria",xlab="Subspecies", ylab="Relative abundance (%)", col = c("#00CC00","#ffff00","#ff3300"))

\# Orbaceae

shapiro.test(data_sub$Orbaceae) #not signifcant, data are normal

#modelo

anova <- aov(Orbaceae ~ Abrev, data=data_sub)

summary(anova)

tukey_ctrl <- LTukey(anova, which="Abrev", conf.level=0.95)

#kruskal.test

kruskal.test(species ~ Orbaceae, data=data_sub)

pairwise.wilcox.test(data_sub $ Orbaceae, data_sub $Abrev, p.adj = "bonferroni", exact=FALSE)

#PLOT

boxplot(Orbaceae ~ Abrev, data=data_sub, cex = 1, main=" Orbaceae ",xlab="Subspecies", ylab="Relative abundance (%)", col = c("#00CC00","#ffff00","#ff3300"))

**Script 3**

**R studio Indic species**

**Protocolo de analise *in silico* do artigo**

**“Africanized honey bees (*Apis mellifera scutellata* x sspp.) gut microbiota: is it close to African or European bees and what it implicates in bee health?”**

install.packages("indicspecies")

library(indicspecies)

library(readr)

library(phyloseq)

packageVersion("phyloseq")

table <- read_csv("indicspecies/table_L6.csv")

View(table)

#indicspeaces

#Subspecies

abund = table[,3:ncol(table)]

subspecies = table$Subspecies

inv = multipatt(abund, subspecies, func = "r.g", control = how(nperm=9999))

summary(inv)

#Site

abund = table[,3:ncol(table)]

Site = table$Site

inv = multipatt(abund, Site, func = "r.g", control = how(nperm=9999))

summary(inv)
